# Bile acid fitness determinants of a *Bacteroides fragilis* isolate from a human pouchitis patient

**DOI:** 10.1101/2023.05.11.540287

**Authors:** Aretha Fiebig, Matthew K. Schnizlein, Selymar Pena-Rivera, Florian Trigodet, Abhishek Anil Dubey, Miette Hennessy, Anindita Basu, Sebastian Pott, Sushila Dalal, David Rubin, Mitchell L. Sogin, A. Murat Eren, Eugene B. Chang, Sean Crosson

## Abstract

*Bacteroides fragilis* comprises 1-5% of the gut microbiota in healthy humans but can expand to >50% of the population in ulcerative colitis (UC) patients experiencing inflammation. The mechanisms underlying such microbial blooms are poorly understood, but the gut of UC patients has physicochemical features that differ from healthy patients and likely impact microbial physiology. For example, levels of the secondary bile acid deoxycholate (DC) are highly reduced in the ileoanal J-pouch of UC colectomy patients. We isolated a *B. fragilis* strain from a UC patient with pouch inflammation (i.e. pouchitis) and developed it as a genetic model system to identify genes and pathways that are regulated by DC and that impact *B. fragilis* fitness in DC and crude bile. Treatment of *B. fragilis* with a physiologically relevant concentration of DC reduced cell growth and remodeled transcription of one-quarter of the genome. DC strongly induced expression of chaperones and select transcriptional regulators and efflux systems and downregulated protein synthesis genes. Using a barcoded collection of ≈50,000 unique insertional mutants, we further defined *B. fragilis* genes that contribute to fitness in media containing DC or crude bile. Genes impacting cell envelope functions including cardiolipin synthesis, cell surface glycosylation, and systems implicated in sodium-dependent bioenergetics were major bile acid fitness factors. As expected, there was limited overlap between transcriptionally regulated genes and genes that impacted fitness in bile when disrupted. Our study provides a genome-scale view of a *B. fragilis* bile response and genetic determinants of its fitness in DC and crude bile.

**Importance:** The Gram-negative bacterium, *Bacteroides fragilis*, is a common member of the human gut microbiota that colonizes multiple host niches and can influence human physiology through a variety of mechanisms. Identification of genes that enable *B. fragilis* to grow across a range of host environments has been impeded in part by the relatively limited genetic tractability of this species. We have developed a high-throughput genetic resource for a *B. fragilis* strain isolated from a UC pouchitis patient. Bile acids limit microbial growth and are altered in abundance in UC pouches, where *B. fragilis* often blooms. Using this resource, we uncovered pathways and processes that impact *B. fragilis* fitness in bile and that may contribute to population expansions during bouts of gut inflammation.

## Introduction

The human gut contains a vibrant community of fungi, protists, archaea, and bacteria. To survive and replicate in this environment, these microbial cells must adapt to complex and varying conditions including gradients in pH, O_2_, nutrients and host-derived compounds such as bile acids that vary longitudinally (i.e., from the stomach to the rectum) as well as transversely (i.e., from mucosa to lumen) (1-8). This physicochemically-dynamic environment influences how microbes interact with each other and their host, and thereby shapes the complex ecological networks of the gut. Studies aimed at defining the mechanisms by which indigenous bacteria adapt to and mitigate chemical stressors encountered in the gut are critical as we work to advance understanding of how bacteria survive disruptions in this ecosystem.

Disruptions to the gut environment as a result of infection, antimicrobial therapy, or surgery can result in blooms of opportunistic microbes (9). A surgical disruption common to patients with severe ulcerative colitis (UC) is ileal pouch anal anastomosis (IPAA), where the terminal ileum is joined to the rectum after colectomy to create a J-pouch. Not surprisingly, IPAA reshapes the physiology of gut, influencing bile acid cycling, water absorption and mucosal physiology (10-14). The microbiome of the nascent ileal pouch adopts a colonic profile with increased numbers of anaerobes that have a shifted metabolism relative to microbes of the ileum (15-17). While this surgical procedure mitigates gut inflammation for some, approximately 50% of UC patients who undergo IPAA eventually develop an inflammatory condition of the ileal pouch, known as pouchitis, which is characterized by symptoms such as rectal bleeding and incontinence (18-20). The etiology of pouchitis remains unclear but several members of the pouch microbiota have been implicated (21-23) including *Bacteroides fragilis*, a Gram-negative, opportunistic bacterium that is commonly isolated from the ileal pouch (23, 24). Due to its ability to thrive in diseased and non-diseased states, *B. fragilis* offers an interesting model for studying bacterial fitness and adaptability to environmental shifts encountered in the mammalian gut. Our goal was to develop a novel cultivar of *B. fragilis* isolated from a UC pouchitis patient to serve as a platform to study genetic factors associated with its fitness in the face of host-derived stressors.

Bile contains a complex mixture of detergent-like bile acids that are secreted into the digestive tract to aid in solubilization of dietary fats. These molecules are a stressor for gut microbes as they can disrupt membranes and cause damage to nucleic acids and proteins (25-28). Primary bile acids produced by the host are typically conjugated to amino acids, which promotes their solubility in an aqueous environment. However, the composition of the bile acid pool in the gut changes after secretion from the common bile duct as microbes hydrolyze conjugated amino acids and chemically modify the steroidal core of primary bile acids to yield secondary bile acids (5). The chemical state of bile acids (e.g., primary vs. secondary; conjugated vs. deconjugated) varies along the digestive tract, drives microbial fitness and shapes microbiota composition (29-34). Moreover, since microbes metabolize bile acids, fluctuations in the microbiota, such as those during gastrointestinal disease, cause shifts in the bile acid pool. For example, an inflamed UC pouch is linked to decreased abundance of secondary bile acids, which typically inhibit growth of gut microbes (22, 35). *B. fragilis* has been described as a ‘bile-tolerant’ species and selective media formulations for *B. fragilis* contain ox bile at a concentration that inhibits most enteric bacteria (36, 37).

As a common low abundance microbe that can bloom in conditions of inflammation, we sought to identify *B. fragilis* genes and gene regulatory responses that affect its interactions with environmental bile acids using two complementary approaches: total RNA sequencing (RNAseq) and randomly barcoded transposon insertion sequencing (RB-TnSeq or BarSeq). We describe the complete genome sequence a *B. fragilis* cultivar from a UC pouchitis patient and its transcriptional response deoxycholate (DC), a secondary bile acid. We further report the first barcoded transposon library in *B. fragilis*, which we used to conduct a genome-wide screen for genetic factors that determine resistance to both deoxycholate and a crude bile extract. This multi-omics investigation provides evidence that survival in the presence of bile acids involves multiple stress mitigation systems. Reduced growth in the presence of deoxycholate correlates with highly reduced expression of protein synthesis machinery which may underlie control of *B. fragilis* populations in healthy individuals. A putative sodium-translocating V-ATPase system, cardiolipin synthase, and select lipoprotein and surface polysaccharide biosynthesis enzymes were identified as critical determinants of fitness in the presence of deoxycholate.

## Results

### Developing a model B. fragilis patient isolate

*B. fragilis* strain P207 was isolated from the J-pouch of a human pouchitis patient (24). This strain bloomed to comprise approximately one-half of the pouch bacterial population within the first year after pouch functionalization (24). Some *B. fragilis* isolates are resistant to multiple antibiotics, which poses challenges during treatment and precludes the use of common genetic tools that rely on antibiotic selection (38). *B. fragilis* P207 was cleared from patient 207 by ciprofloxacin treatment (24), and is sensitive to tetracycline, erythromycin and chloramphenicol *in vitro*, which facilitated the use of classical tools for genetic manipulation. *B. fragilis* P207 has other features that make it amenable to *in vitro* study including robust growth and increased aerotolerance compared to *B. fragilis* strains from other pouchitis patients in our collection (24), or to other commonly studied *B. fragilis* strains such as NCTC 9343 and 638R.

To facilitate the development of *B. fragilis* P207 as a genetic model system, we sequenced genomic DNA using a combination of long- and short-read approaches and assembled the reads *de novo* using the repeat graph assembly algorithm of Flye (39), followed by an assembly polishing step in Pilon (40). The complete, circular P207 genome is 5.04 Mbp; no episomal sequences were present in the final assembly. Automated annotation using the prokaryotic genome annotation pipeline (PGAP) (41) predicted a total of 4,110 genes, with 6 ribosomal RNA loci, and one type IIC CRISPR system. *B. fragilis* P207 lacks sequence related to the *Bacteroides* pathogenicity island (BfPAI) (42) and thus does not contain known forms of the *bft* gene (43-45), which encodes the fragilysin toxin. *B. fragilis* P207 is therefore classified as a non-toxigenic *B. fragilis* (NTBF) strain.

#### Characterization of *B. fragilis* growth in bile

To investigate physiological responses of *B. fragilis* P207 to bile, we first sought to identify concentrations of purified bile salts and crude bile extract that attenuate, but do not completely inhibit growth. We cultivated *B. fragilis* P207 in supplemented brain heart infusion medium (BHIS) containing increasing concentrations of the secondary bile acid, deoxycholate (DC), or bile salt mixture (BSM; a 1:1 mixture of deoxycholate and cholate), and measured the culture density after 24 hours. Adding 0.01% (w/v) BSM or DC resulted in a 20 and 30% reduction in the terminal density of *B. fragilis* P207 cultures, respectively (Figure 1). A 0.01% DC concentration is congruent with levels measured in the colon of healthy humans (46) and in uninflamed pouches of FAP patients (35). We similarly examined the effect of crude bile extract from porcine (BEP) on growth of *B. fragilis* P207. In the presence of higher concentrations of crude BEP, growth was delayed (Figure S1) and cultures required a second day to reach saturation (Figure 1). Nevertheless, final culture densities were enhanced by elevated BEP suggesting that after adaptation, P207 can utilize components of crude bile to support growth. In the presence of elevated concentrations of crude bile, cells grew in a biofilm-like aggregate at the bottom of the culture tube, consistent with bile-stimulated biofilm development observed in *Bacteroides* species (47, 48).

**Figure 1:**
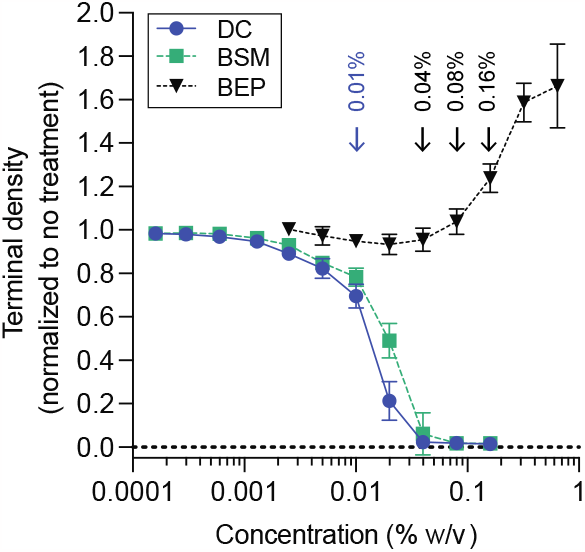
*B. fragilis* P207 growth in the presence of increasing concentrations deoxycholate (DC), bile salt mixture (BSM) or bile extract of porcine (BEP). Data reflect the terminal density (OD_600_) of 3 ml cultures after 24 hours growth for DC and BSM or 48 hours for BEP; mean ± SD of 8 independent trials each normalized to an untreated control. Representative growth curves for each condition are presented in Figure S1. Blue arrow highlights the concentration of DC and BSM used for subsequent experiments. Black arrows highlight the concentrations of BEP used for subsequent experiments.

#### Deoxycholate treatment remodels the *B. fragilis* transcriptome

To identify *B. fragilis* genes that function in adaptation to bile acid exposure, we first conducted an RNA sequencing experiment to identify transcripts that have altered abundance upon treatment with 0.01% DC. This physiologically relevant concentration of DC, which modestly reduces growth of *B. fragilis* P207 (Figure 1), induced rapid and global changes in the transcriptome. Within 6 minutes of exposure approximately 10% of *B. fragilis* transcripts (427 / 4110) had significantly altered abundance (|log_2_(fold change)| > 1.5 and FDR p < 1 E^-10^); after 20 minutes of DC exposure the fraction of regulated genes meeting this threshold was over 25% of the genome (1071 / 4110) (Table S1, Figure 2A-B). As expected, the transcriptional response at 6 and 20 minutes was highly correlated with the magnitude of change increasing as a function of exposure time (Figure 2C). We briefly summarize our transcriptome analyses here and provide a more thorough discussion of this complex data set in the supplemental material (**Supplemental Results**). Interpro (49) and GO term (50) assignment followed by gene set enrichment analysis (GSEA) (51), and Fisher’s exact test (52) revealed gene function classes that are significantly (FDR p <0.05) influenced by DC treatment (Table S2). Functional categories associated with acute stress response and mitigation of protein misfolding were significantly enriched in upregulated genes (Table S2), suggesting DC induces a severe stress response. Functional categories associated with protein translation were significantly enriched in down regulated genes (Table S2), which is consistent with a metabolic shift toward slower growth. Additionally, DC triggered dysregulation of cellular ion homeostasis and modulation of expression of efflux systems known for exporting toxic compounds.

**Figure 2:**
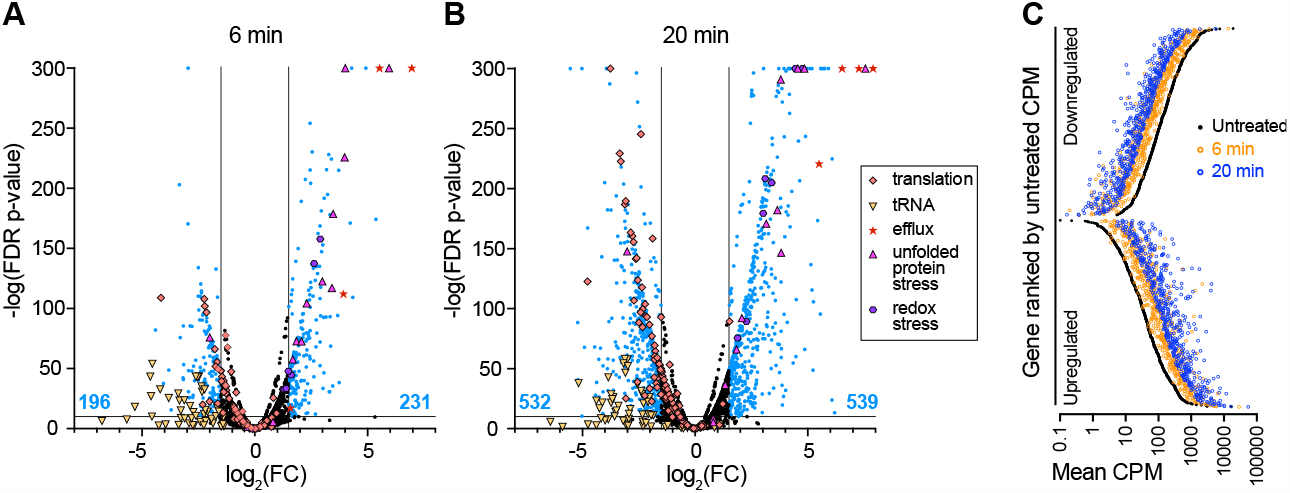
Treatment of *B. fragilis* P207 with a sub-lethal concentration of deoxycholate (DC) induces large-scale activation and repression of transcription. Volcano plots of differentially transcribed genes at **(A)** 6 minutes and **(B)** 20 minutes post DC exposure. Lines indicate cutoff criteria (FDR p < 10^−10^ and absolute log_2_(fold change) >1.5, where fold change (FC) reflects the ratio of CPM after DC exposure / CPM before DC exposure). The number of up- or down-regulated transcripts (blue points) at each time point is indicated at the bottom of graph (blue text). Genes that do not meet the cutoff criteria are in black. Functional categories of genes are highlighted with special symbols include: a) translation processes (GO terms: 0003735, 0006414, 0006400, 0043022, 0000049; orange diamonds), b) transmembrane efflux processes (operon *PTOS_003611-3614;* red stars), c) unfolded protein stress (GO term: 0051082; pink triangles), d) redox stress (catalase, *msrB*, and *dps*; purple hexagons), e) tRNA (related to translation processes; yellow inverted triangles). A subset of transcriptional changes were confirmed with RT-qPCR (Figure S2). **C**. Differentially transcribed genes 6 minutes (orange circles) and 20 minutes (blue circles) after 0.01% (w/v) DC exposure are highly correlated. The 1071 genes that are significantly up- and downregulated by 20 minutes post DC treatment relative to untreated control (black circles) are ranked by transcript counts per million (CPM) of gene expression in the untreated condition.

The global transcriptional response to DC exposure is consistent with large-scale remodeling of the physiological status of the cell to slow growth, shift carbon and energy metabolism, and to mitigate a constellation of cellular stresses that arise due to bile exposure. Although these transcriptomic data are informative, the presence of over 1000 regulated transcripts posed challenges in assigning the contribution of specific genes or pathways to *B. fragilis* fitness in bile. As such, we sought to extend our transcriptomic study by implementing a genome-scale genetic approach to directly identify genes that contribute to *B. fragilis* P207 fitness in the presence of DC and a crude bile extract.

### Transposon mutagenesis of B. fragilis P207 defines a set of candidate essential genes

To enable genetic analysis of *B. fragilis* P207 physiology, including bile resistance, we generated a pool of >50,000 barcoded Tn-*Himar* insertion mutants for random barcode transposon insertion site sequencing (RB-TNSeq or BarSeq) (53, 54). Mapping the insertion sites in the pool identified 49,543 reliably mapped barcodes at 43,295 distinct sites in the genome. This *B. fragilis* Tn-*Himar* pool contains a median of 8 mutant strains per gene, and at least one insertion in the central region of 87.9% of protein coding genes (Table S3). In parallel, we produced a collection of 889 Tn-*Himar* mutant strains that were arrayed and individually mapped (Table S4). This arrayed collection contains strains carrying transposon insertions in approximately 14% of the predicted genes in the *B. fragilis* P207 genome.

We employed a probabilistic approach that relies on hidden Markov models (HMMs) to quantify gene essentiality in mutant libraries (55), and identified 310 candidate essential genes and 258 genes for which Tn-*Himar* insertions are predicted to yield growth defects in BHIS medium (Table S5). We further used a Bayesian (Gumbel) approach (56) to assess gene essentiality (Table S5). As less than one-quarter of TA dinucleotide sites carry insertions in the pool, these candidate essential lists are likely incomplete. Nonetheless, expected essential genes including those with key roles in cell division, cell envelope biogenesis, DNA replication, transcription, and translation were common to both the Gumbel and HMM essential lists. Additional data and discussion of candidate essential genes in *B. fragilis* P207 is presented in the **Supplemental Results**.

#### Barcoded transposon mutagenesis identifies genes that determine *B. fragilis* fitness in bile

To identify *B. fragilis* genes impacting fitness in bile through barcoded transposon sequencing (BarSeq), we used treatment dosages that moderately inhibited growth. This approach allowed for concurrent identification of Tn-*Himar* mutant strains both more susceptible and more resistant to treatment. We cultivated the barcoded Tn-*Himar* mutant pool in BHIS broth with either 0.01% DC, 0.01% BSM, or a range of BEP concentrations (0.04, 0.08 and 0.16%). In parallel, we cultivated the pool in plain BHIS medium to differentiate strains with general growth defects from those with bile specific growth defects. All cultures were serially passaged twice to amplify fitness differences between mutant strains; barcode abundances were evaluated after the first and second passages. Fitness scores for each gene in each condition were calculated using the approach of Wetmore *et al*. (54) and are presented in Table S6. Briefly, fitness scores represent the composite fitness advantage or disadvantage of all strains bearing insertions in a particular gene relative to a control condition. Negative fitness scores indicate mutants that are impaired, and hence genes that support fitness in the presence of bile. Positive fitness scores indicate mutants that are advantaged in bile; the presence of these genes is therefore detrimental in bile. Statistical t-scores – which take into account variance in fitness of each transposon insertion strain for each gene (54) – are presented in Table S6.

Principal component analysis of the genome-scale fitness data showed high experimental reproducibility between biological replicates (Figure S3). The fitness profiles of the *B. fragilis* P207 mutant pool cultivated in DC and BSM were more similar to each other than to crude BEP, which was expected given that BSM is a purified mixture of cholate and DC while BEP is a biochemically complex animal extract. Exposure to DC had a greater fitness impact on *B. fragilis* than exposure to an equivalent concentration of BSM. To validate the Barseq fitness measurements, we selected clones from our arrayed collection of individually mapped mutants (Table S4) that harbored insertions in genes identified as fitness factors by BarSeq. These mutants displayed defects in growth rate, terminal density, and/or lag time in the presence of DC (Figure S4). Growth of these individual mutant strains was consistent with the fitness scores for the corresponding genes derived from BarSeq, lending confidence to the genome scale data set.

Genes that impacted fitness in each bile condition were defined as those with average fitness scores less than -4 or greater than +1.5 after the second passage, excluding genes that affected growth in plain BHIS medium. Using these criteria, 14 genes were identified as determinants of *B. fragilis* P207 fitness in 0.01% BSM, 63 in 0.01% DC, and 89 in at least one concentration of BEP. Together this represents a set of 122 genes that significantly influence growth in at least one bile treatment condition. Clustering this gene set based on fitness scores revealed high overlap in groups of genes that positively or negatively contributed to *B. fragilis* fitness across all conditions, and further revealed a set of genes that contributed to fitness in a specific bile condition (Figure 3). No gene was identified as a BSM-specific fitness factor. As will be discussed below, genes that function in membrane transport, stress responses, cell envelope biosynthesis and energy metabolism are among the major bile fitness factors.

**Figure 3:**
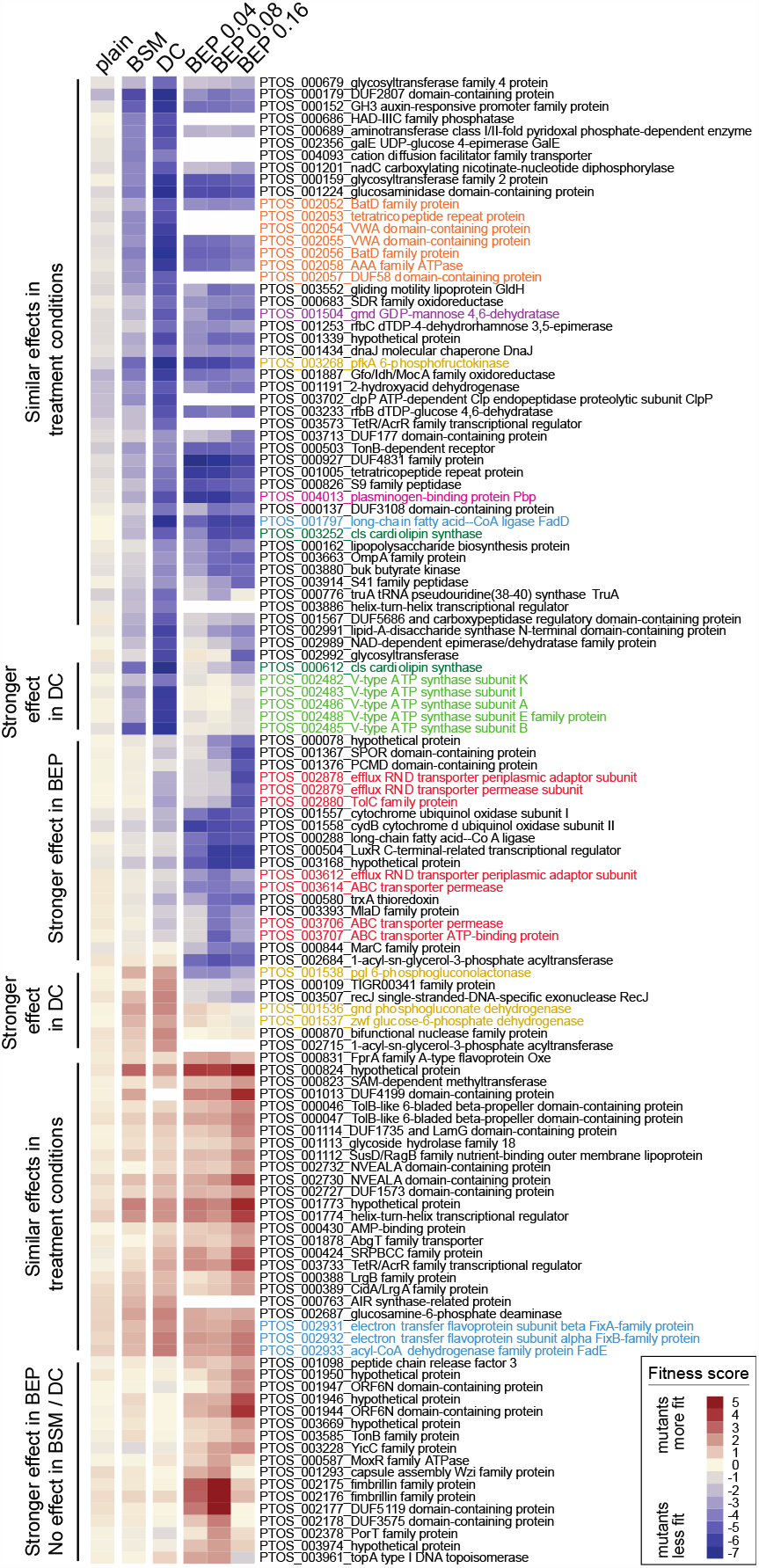
Genes that contribute to *B. fragilis* fitness in the presence of purified bile acids and crude bile have diverse metabolic, stress response, and cell envelope functions. Heat map of fitness scores for the 122 genes that are significant determinants of *B. fragilis* fitness in at least one treatment condition (see Materials and Methods for significance thresholds); treatment conditions are arranged in columns and genes in rows. Composite fitness scores for genetic mutants grown in plain BHIS medium without bile are labeled (plain). Mutant strain growth was measured in BHIS containing 0.01% (w/v) bile salt mixture (BSM), 0.01% (w/v) deoxycholate (DC), and 0.04%, 0.08%, and 0.16% bile extract of porcine (BEP). Gene-level fitness scores were hierarchically clustered; gene arrangement was manually adjusted to group genes presumed to be in operons. White blocks indicate genes with insufficient barcode counts in the reference condition of a particular experiment to calculate a fitness score. Colored gene names highlight select functional categories discussed in the text: red – efflux systems, light green – V-ATPase operon, dark green – cardiolipin synthase genes, light blue – lipid metabolism, mustard – central carbon metabolism, orange – Bat aerotolerance operon, fuchsia – plasminogen-binding protein (Pbp), and purple – *gmd* protein glycosylase.

#### Transcriptional regulation does not predict fitness contribution

Most transcriptionally regulated genes have neutral or modest fitness scores in DC when disrupted. Moreover, transcription of genes with highly significant fitness scores was largely unaffected by DC treatment (Figure 4). The small number of genes that meet our significance thresholds for both transcriptional regulation and fitness in DC are presented in Figure S5. While the difference in gene sets revealed by Tn-seq and BarSeq analysis of *B. fragilis* responses to bile acid may initially seem surprising, observing a change in transcription in a particular condition does not necessarily mean that the change is functionally relevant (57). This was evident in early genome-scale genetic analyses in prokaryotes and eukaryotes (58, 59) as well as studies that have integrated transcriptomic and functional genomic analysis (60, 61). Each show that that gene disruption often did not lead to the phenotypic changes that would be expected based on transcriptional regulatory profiles.

**Figure 4:**
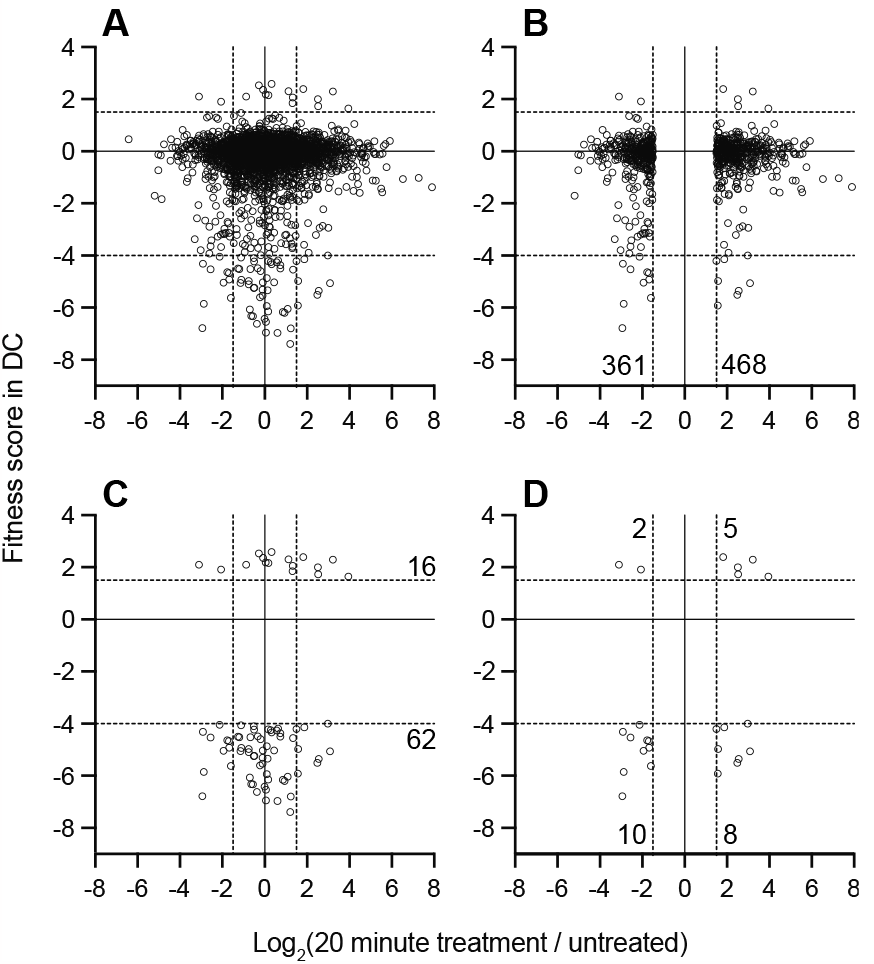
Genes that are transcriptionally regulated by DC treatment are weakly correlated with genes that determine fitness in medium containing DC. Log_2_(fold change) in transcript levels before and after 20 minutes of exposure to 0.01% DC (x-axis) versus gene-level fitness scores after cultivation in 0.01% DC (y-axis) for the 3270 genes with fitness scores. Each point is a gene, for **(A)** all genes with fitness scores, **(B)** transcriptionally regulated genes with fitness data, **(C)** genes that pass fitness criteria, or **(D)** transcriptionally regulated genes that also pass fitness criteria. The 25 genes in **D** are listed in Figure S5. Dotted lines indicate cutoff thresholds on each axis. Numbers on graph indicate the number of genes in each region of the graph that meet the selection criteria.

Though there is little overlap in the specific genes that are transcriptionally regulated and those that have fitness effects when disrupted (Figure 4), the functional classes of genes identified in these complementary studies reveal a consistent physiological response. Both data sets provide evidence for the importance of efflux and stress response systems in response to bile exposure; we provide an in-depth presentation and discussion of the genes contained in these two categories in the **Supplemental Results**. Moreover, both experimental approaches provide evidence for physiological shifts consistent with slower growth, energy limitation, and envelope remodeling during bile challenge. Of the 825 genes lacking fitness data, nearly 30% exhibit changes in transcript levels 20 minutes after DC exposure, a proportion similar to the rest of the genome. Certainly, transcriptional regulation of some essential genes may be important for *B. fragilis* fitness in DC; such genes are missed by BarSeq analysis. Using these complementary genome-scale methods offers a more complete understanding of *B. fragilis* physiology when challenged with bile acids.

#### Barseq identifies metabolic branchpoints that impact bile resistance

Genes that function at select metabolic branchpoints had some of the most positive and most negative fitness scores in our BarSeq data set and provide insight into the metabolic constraints imposed by bile. Glucose-6-phosphate (G6P) connects major pathways in central carbon metabolism, and can be shunted towards glycolysis to generate ATP and NADH, or to the pentose phosphate pathway to yield NADPH and pentose sugars (Figure 5A). Disruption of phosphofructokinase (*pfkA*; *ptos_003268*), which catalyzes the committed step in directing G6P toward glycolysis, resulted in severe fitness defects in all bile treatment conditions (Figure 3 & 5, Table S6). In contrast, strains harboring insertions in genes comprising the oxidative branch of the pentose phosphate pathway (*zwf, pgl* or *gnd*; *ptos_001536-38*) had a fitness advantage in the presence of DC (Figures 3 & 5, Table S6). From these results, we infer that in the presence of DC an active glycolytic pathway is more advantageous than shunting glucose to the pentose phosphate pathway. *pfkA* is constitutively expressed, but the *gnd*-*zwf*-*pgl* operon is transcriptionally activated by acute DC exposure (Figure 5C, Table S1).

**Figure 5:**
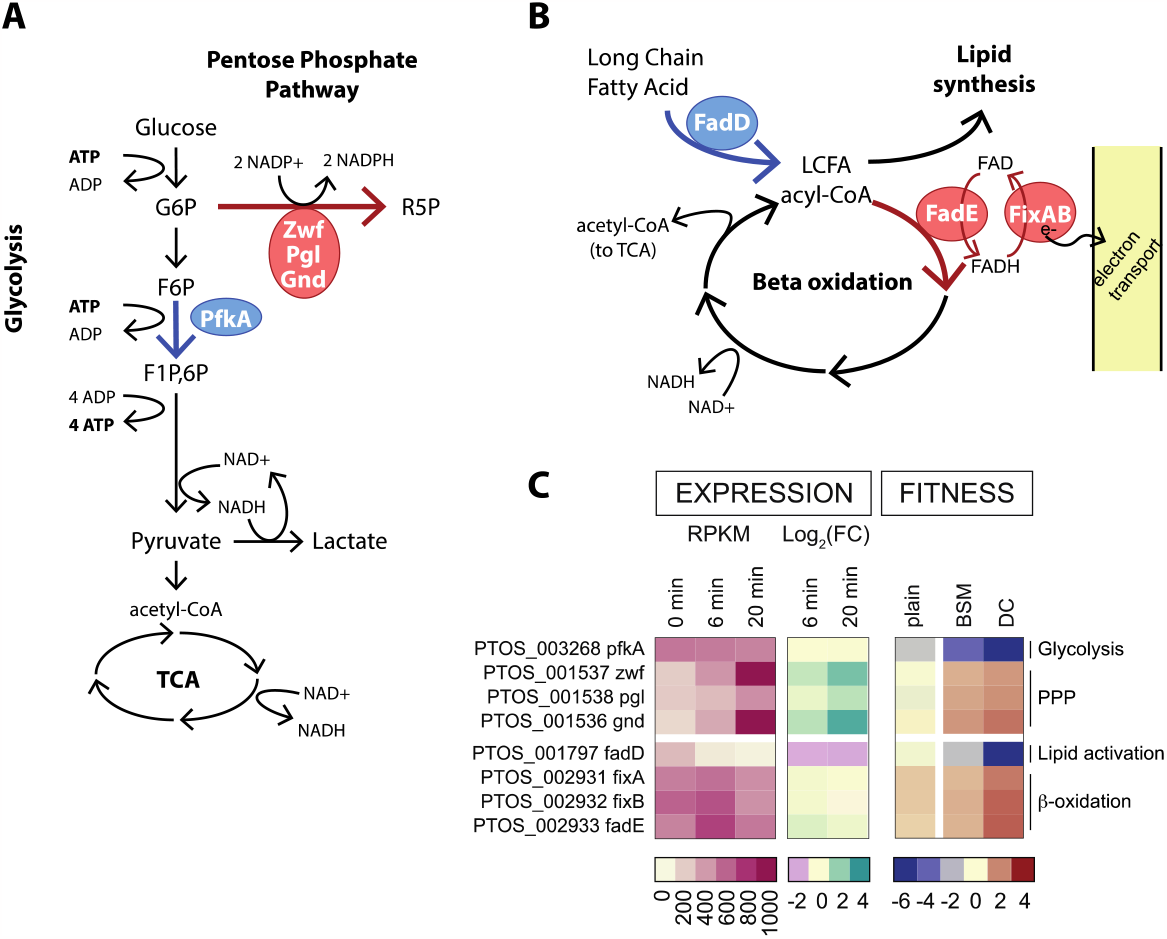
Fitness scores highlight metabolic branchpoints in central carbon and lipid metabolism. **A**. Schematic of glucose metabolism pathways highlighting the branchpoint between glycolysis and the pentose phosphate pathway. **B**. Schematic of lipid metabolism highlighting the fate of long-chain fatty acids (LCFA). Blue and red arrows and enzyme circles highlight processes that promote or inhibit fitness in the presence of DC, respectively. Pathway names are bold, key enzymes are named in circles, metabolites are in plain text. **C**. Heat map of expression values and fitness scores for the genes highlighted in A and B. Transcript abundance is presented as reads per kilobase per million reads (RPKM) and regulation is indicated by the log_2_(fold change) compared to untreated cells (0 min). Color scales representing absolute expression, fold change in expression, and mutant fitness scores are presented below each type of data.

Genes involved in fatty acid metabolism also exhibited contrasting fitness scores. Disruption of long chain fatty acid-CoA ligase (*fadD; ptos_001797*), which activates long-chain fatty acids for either anabolic processes or beta-oxidation, resulted in extreme sensitivity to all treatment conditions (Figure 3 & 5, Table S6). The first step in beta oxidation is executed by the products of *fadE* and *fixAB* (*ptos_002931-33*), and disruption of these genes conferred a fitness advantage in the tested conditions (Figure 3 & 5). These results support a model in which activation of fatty acids by *fadD* for anabolic, rather than catabolic processes, is important for bile acid resistance. Although *fadD* is critical for fitness, transcription of all three LCFA-CoA ligases in *B. fragilis* is reduced upon DC exposure providing an example of incongruence in direction of transcriptional regulation and expected fitness impact. Transcription of *fadE-fixAB* is constitutive in all conditions tested (Figure 5, Table S2). We note that beta-oxidation and the oxidative branch of the pentose phosphate pathway generate reducing equivalents (FADH or NADPH), and disruption of genes in either of these pathways is advantageous in the presence of bile.

#### The contribution of membrane bioenergetic systems to fitness in bile

*B. fragilis* P207 encodes two distinct ATP synthase systems: a standard F-type (F_0_F_1_) system (PTOS_001821-29) that utilizes a proton motive force to generate ATP, and a second V-type ATPase (PTOS_002482-88) that has all the conserved residues to coordinate and translocate sodium ions (Figure 6, (62-64)). We recovered few strains with transposon insertions in the genes encoding the F_0_F_1_ system in our mutant pool and were therefore unable to evaluate the fitness of F_0_F_1_ mutants. These genes are transcriptionally upregulated by DC exposure, a result that suggests energy limitation in the presence of bile (Figure 6, Table S1 & S2). On the other hand, strains with transposon insertions in the genes encoding the V-type system were well represented in the pool, but were nearly undetectable after cultivation in DC. The fitness defects of V-type ATPase mutants were among the most extreme in the entire DC dataset (Figure 3, Table S6), and these defects were largely specific to DC.

**Figure 6:**
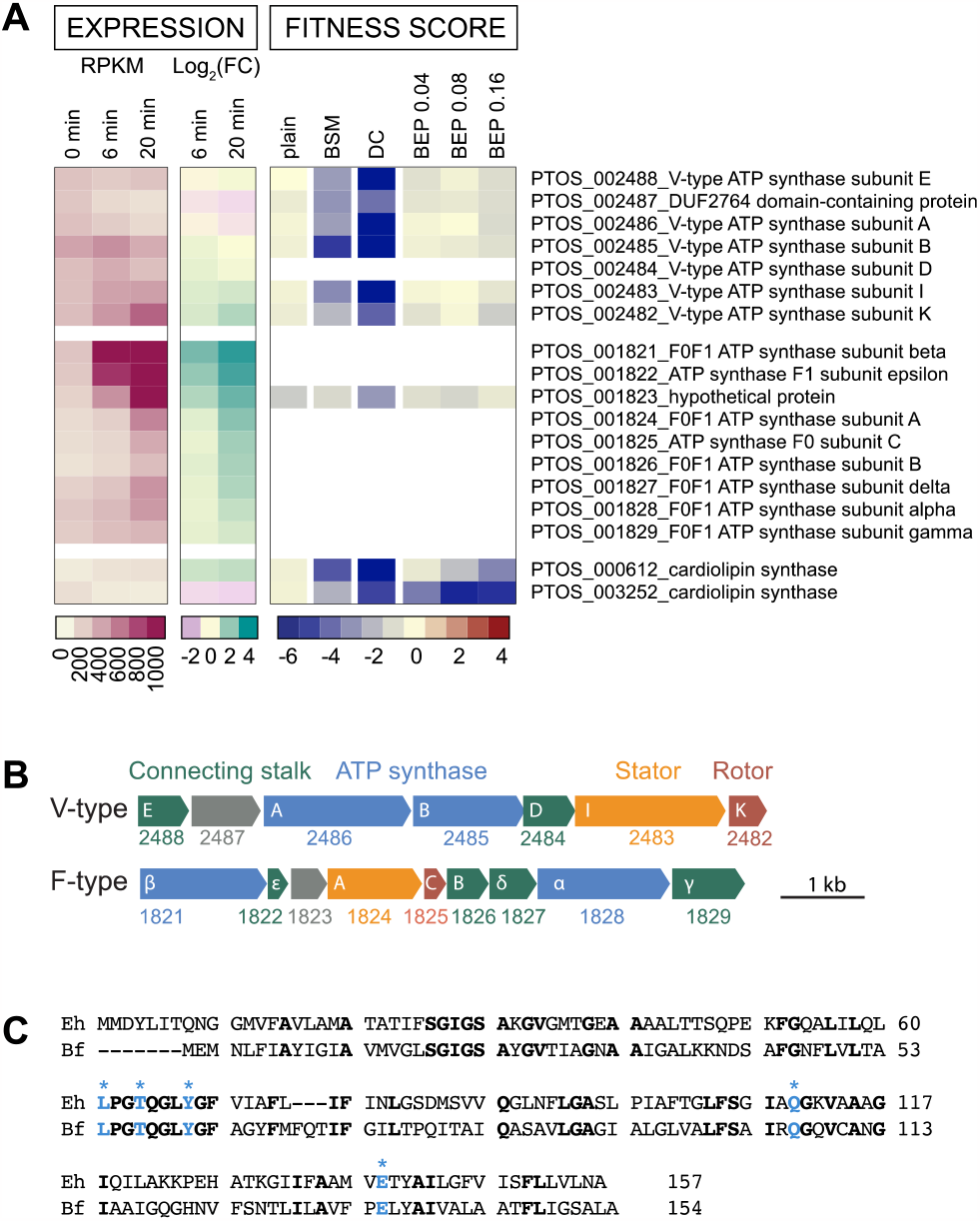
B. *fragilis* encodes two rotary ATP synthase / ATPase systems; the V-type system bears conserved residues for sodium ion translocation and is a critical fitness factor in deoxycholate. **A**. Expression level and fitness scores for genes in the V-type and F-type ATP synthase / ATPase operons, presented as in Figure 5. White blocks indicate genes with insufficient barcode counts in the reference condition to calculate a fitness score. **B**. Operon structure of F-type and V-type ATPase systems. PTOS gene locus numbers are below gene outlines; annotated subunit names are in white. Genes are colored as follows: cytoplasmic ATP synthesis/hydrolysis subunits – blue; transmembrane, ion-translocating, rotary subunits – red; transmembrane stator subunits – orange; stalk subunits connecting the enzymatic complex to the membrane complex – green; and unknown function - grey. The V-type system has fewer connecting accessory subunits than the F-type system or other characterized bacterial V-type systems (95). **C**. Protein sequence alignment of the K subunits of *B. fragilis* (Bf) and the sodium ion translocating *Enterococcus hirae* (Eh) V-ATPases. Identical residues are bold. Residues experimentally determined to coordinate sodium ions in *E. hirae* (64) are highlighted in blue with asterisk.

#### Cell envelope features contribute to bile resistance

Bile salts are detergent-like molecules that can disrupt membranes, so it was expected that genes involved in biosynthesis of components of the *B. fragilis* cell envelope would impact fitness in the presence of bile. The phospholipid, cardiolipin, has been implicated in adaptation and/or resistance of bacteria to envelope stress conditions including high osmolarity (65) and bile stress (61, 66). *B. fragilis* encodes two predicted cardiolipin synthases, *ptos_000612* and *ptos_003252*, and disruption of either of these genes resulted in sensitivity to bile. Thus, the functions of these genes are not entirely redundant under the tested conditions (Figure 6, Table S6). Both cardiolipin synthase genes are modestly, and oppositely, regulated at the transcriptional level (Figure 6, Table S1).

Many proteins and lipids of the *B. fragilis* cell envelope are heavily glycosylated. *gmd* (*ptos _001504;* GDP-mannose 4,6-dehydratase) and *fcl* (*ptos_001503*; GDP-L-fucose synthase) function in protein O-glycosylation, which is important for host colonization (67, 68). Transposon insertions in *gmd* resulted in a severe fitness defect when *B. fragilis* P207 was cultivated in the presence of DC and crude bile extract. Disrupting GDP-L-fucose synthase (*ptos_001503, fcl)* also reduced fitness in bile, though the impact was less severe (Figure 3, Table S6). A target of the *gmd-fcl* glycosylation pathway is a glycosyltransferase encoded by *ptos_002992* (69), and strains with insertions in this gene also had reduced fitness in bile. Several other genes with predicted functions in biosynthesis of capsular polysaccharide (CPS), lipopolysaccharide (LPS) or other surface polysaccharides (e.g. *ptos_000679, ptos_000683, ptos_000686, ptos_000689, ptos_000159, ptos_000162, ptos_001253, ptos_001293, ptos_002991, ptos_002992, ptos_003233*) also resulted in sensitivity to bile treatments. Among characterized *B. fragilis* surface-exposed lipoproteins (70, 71), the highly-expressed plasminogen-binding protein (Pbp; *ptos_004013*; (70)) conferred resistance across all conditions and was among the most important DC resistance factors in our dataset (Figure 3, Table S6). This result is consistent with data from *B. thetaiotaomicron* showing that Pbp supports fitness in the presence of a range of bile compounds (53).

#### Specific fitness factors in crude porcine bile

While our Barseq study yielded few specific hits for BEP, we identified genes connected to LPS export and several genes of unknown function that are linked to fitness in BEP. Insertions in two efflux systems (*ptos_002878-80* and *ptos_003611-14*) and in components of the predicted lipopolysaccharide export system (*ptos_003706-07*) were more detrimental to *B. fragilis* fitness in BEP than in DC (Figures 3 & S7). Additionally, strains with disruptions in a cluster of genes of unknown function (*ptos_001944, ptos_001946, ptos_001947* and *ptos_001950*) and two genes encoding fimbrillin family proteins (*ptos_002175-76*) resulted in a specific fitness advantage in crude bile.

## Discussion

### A new resource for the study of *B. fragilis* physiology

We have developed *B. fragilis* isolate P207 as a new model system to investigate genes, metabolic pathways, and cellular processes that contribute to *B. fragilis* fitness in conditions encountered in the inflamed mammalian gut. Other well-studied strains of *B. fragilis* have been isolated from infection sites where this species was an opportunistic pathogen (e.g. NCTC 9343 from an appendix abscess, and 638R from an abdominal abscess). Strain P207 was isolated from the distal bowel of an ulcerative colitis patient who experienced inflammation of the ileal-rectal pouch, or pouchitis (24). Using this strain we generated a diverse pool of barcoded Tn-*Himar* mutants, which can be used to interrogate strain-level fitness at the genome scale in any condition of interest. To our knowledge, this is the first barcoded transposon mutant library in *B. fragilis*, a species that is typically less amenable to molecular genetic analysis than other well-studied *Bacteroides* species. We have further generated an arrayed collection of individually mapped transposon insertion mutants that captures ≈14% of genes in the genome and that can be used for directed studies of individual *B. fragilis* genes.

Bile acids play a major role in shaping the gut microbiota and bile acid profiles are reported to shift in the inflamed gut. Specifically, UC inflammation is associated with increases in primary bile acids and decreases in secondary bile acids such as deoxycholate (DC) (35). DC concentrations that attenuated growth of *B. fragilis* P207 in vitro are similar to those found in healthy individuals. DC was undetectable in the stool of patient 207 (72) when *B. fragilis* P207 bloomed to become the dominant species in the pouch (24). Thus it is reasonable to conclude that the absence of deoxycholate contributed, at least in part, to the population expansion of *B. fragilis* in patient 207, though other factors relating to the overall metabolic versatility of *B. fragilis* (73, 74) likely contributed to its bloom as well.

As a potent and relevant inhibitor of *B. fragilis* growth, we sought to understand the genetic factors that enable *B. fragilis* to survive bile acid challenge using two complementary genome-scale approaches. The BarSeq and RNA-seq approaches that we report here identified largely distinct sets of genes, which collectively inform the physiological underpinnings of *B. fragilis* bile tolerance. RNA-seq revealed massive transcriptional reprograming upon exposure to a physiologically relevant concentration of DC. This data set evidenced a physiologic shift involving a reduction of protein synthesis capacity and enhanced stress mitigation processes (Figure 2, Table S2 & Supplemental Results), which is consistent with observations from other enteric microbes for which bile responses have been studied (26). BarSeq identified a smaller set of genes that specifically contributed to fitness in our bile conditions, many of which are constitutively expressed. As discussed below, many of the major insights from the fitness data (the conditional requirements a sodium translocating alternative ATPase and two cardiolipin synthases, and identification of key metabolic branchpoints) support a model in which deoxycholate severely compromises the cell envelope, the ability of the cell to maintain a proton motive force, and the ability of the cell to generate ATP.

### Cardiolipin and bile acid resistance

The anionic phospholipid, cardiolipin, is not typically an essential component of bacterial membranes but plays a critical role in supporting function of trans-membrane protein complexes including respiratory complexes and ATP synthase (75-79). *B. fragilis* contains two cardiolipin synthase genes, *ptos_000612* and *ptos_003252*, which both contribute to fitness in the presence of bile. Several lines of evidence indicate that cardiolipin stabilizes bacterial membranes in the presence of bile acids. For example, fitness defects have been reported for cardiolipin synthase mutants in *Entercoccus faecium* exposed to bile (61), and elevated levels of cardiolipin have been reported in *Lactobacillus* adapted to growth in sub-lethal concentrations of bile salts (66). In addition, phospholipid vesicles containing high fractions of cardiolipin are more resistant to solubilization by bile salts than vesicles with low cardiolipin (66).

In addition to facilitating the dynamic movement of membrane protein complexes, cardiolipin acts as a proton trap to facilitate transfer of protons exported during electron transport to ATP synthase (77, 80-82). This role as a proton trap is important in the case of bile acid exposure because membrane stresses are associated with increased proton leakage, and dissipation of proton motive forces (PMF). Higher local concentrations around the ATP synthase provided by CL would help maintain ATP synthesis. Indeed, exposure to membrane decoupling agents leads to elevated levels of cardiolipin (83), providing evidence that this lipid may be regulated to mitigate loss of membrane potential during stress. DC degenerates PMF at concentrations found in the distal bowel of healthy patients, which are similar to the DC concentrations used in this study (27, 35). The collapse of proton gradients across the bacterial cell membrane compromises the function of ATP synthase(s), which can both harness the potential energy of trans-membrane ion gradients to synthesize ATP or use the chemical energy stored in ATP to pump ions across the membrane against a chemical gradient (depending on the concentrations of ATP and ions).

#### The contribution of membrane bioenergetic systems to fitness in bile

The fitness profiles of the two *Bacteroides* ATP synthase systems provide insight into the distinct contributions of these systems under different conditions. While the F_0_F_1_ system is a critical fitness determinant in standard cultivation conditions, the V-type system is dispensable under standard conditions, but essential in the presence of DC. In eukaryotic cells, V-type ATPases have been characterized as proton pumps that function to acidify vacuoles, but V-type systems in bacteria (also called A-type) can couple ion motive forces to ATP synthesis (e.g. *Thermus thermophilus* (84)), or expend ATP to pump ions from the cell to maintain homeostasis (e.g. in *Enterococcus hirae* (85)). The function of the V-type system in *Bacteroides* is not known. Considering ***a)*** the detrimental effect of bile acids on membrane integrity, ***b)*** the presence of conserved Na^+^-coordinating residues in the system (Figure 6), and ***c)*** the fact that membranes are less permeable to Na^+^ than H^+^ (86), we hypothesize that the V-type system may function primarily as an ATP synthase that harnesses sodium motive force to support ATP production when membrane disruptors such as DC are present. ATP synthases that can translocate Na^+^ rather than H^+^ are potentially advantageous in conditions where proton gradients can become compromised (62).

The operon encoding the V-type ATP synthase is broadly conserved in the phylum Bacteroidetes including in *Porphyromonas gingivalis* where it is reported to be upregulated in the presence of sapeinic acid, a host-derived lipid that disrupts bacterial membranes (87). Expressing multiple ATPase/synthase systems likely enables bacteria to leverage distinct ion gradients to support fitness in niches where proton gradients may be unreliable due to extreme pH, high temperature or an abundance of membrane disrupting chemicals (62). Notably, *B. fragilis* encodes a sodium pumping oxidoreductase, Nqr, that facilitates the establishment and maintenance of Na^+^ gradients during electron transport; Nqr accounts for about 65% of the NADH:quinone oxidoreductase activity of the cell (88) indicating that the maintenance of a sodium gradient is an important aspect of *B. fragilis* physiology. Sodium concentrations can become elevated in the inflamed the gut of UC patients (89), and bacteria that can leverage elevated sodium levels for energy production may have an growth advantage.

To our knowledge the *Bacteroides* V-type ATPase/synthase has not been functionally characterized. It has features of other V-type systems - such as a duplicated c/K subunit that forms the ion-conducting pore - but has a distinct number of subunits compared to characterized ATPase/synthase operons. In addition to the cytoplasmic alpha and beta subunits that form the catalytic domain, the ion translocating k subunit, and the stator subunit I, the *Bacteroides* V-type operon encodes only three additional proteins to coordinate ion translocation with ATP synthesis/hydrolysis (Figure 6), which is fewer than other bacterial V-type ATPase systems (90). The second gene in the *Bacteroides* V-ATPase operon, *ptos_002487*, is annotated as DUF2764. This conserved protein is exclusively bacterial and most commonly found in the phylum Bacteroidetes (49). The genes comprising this unique V-ATPase genetic element are clearly important fitness determinants in the secondary bile acid, DC, and merit further study.

## Materials and Methods

### Bacterial strains and primers

Strains and primers used in this study are listed in **Table S7**. All primers were synthesized by Integrated DNA Technologies (Coralville, IA, USA).

### Growth media

*B. fragilis* strain P207 was grown in BHIS medium (37 g/L Bacto™ Brain Heart Infusion (Becton, Dickinson and Company), 10 g/L Yeast Extract (Fisher BioReagents), 0.5 g/L L-cysteine (Sigma), supplemented after autoclaving with 5 ug/ml hemin and 1 ug/ml vitamin K. *B. fragilis* is resistant to gentamicin and in some cases, gentamicin (20 ug/ml) was added to prevent contamination or counter select *E. coli*. Erythromycin (5 ug/ml) was added to select transposon bearing strains. Solid BHIS plates contained 1.5% agar (Lab Scientific, A466) and 0.001% EDTA. *B. fragilis* manipulations were done aerobically on the benchtop. Incubations were carried out at 37°C in an anaerobic chamber (Coy Laboratory Products, Grass Lake, MI) filled with 2.5% hydrogen, 97.5% nitrogen. *Escherichia coli* strain AMD776 was grown in LB broth (1% peptone, 0.5% yeast extract, 1% NaCl) supplemented with 100 ug/ml carbenicillin and 0.3 mM diaminopimelic acid (LB-Carb-DAP). *E. coli* was grown aerobically at 37°C unless otherwise noted.

### Dose response of B. fragilis *P207 to bile salts*

Starter cultures of *B. fragilis* P207 were inoculated from freezer stocks into BHIS. After overnight growth, these cultures were diluted to a starting OD600 of 0.01 in 3 mL of BHIS supplemented with increasing concentrations of 1) bile salt mixture (50% cholate and 50% deoxycholate; Sigma-Aldrich B8756), 2) deoxycholate (Fisher Scientific BP349), or 3) porcine bile extract (Sigma-Aldrich B8631). Cultures were incubated anaerobically in 14 mm glass tubes. Optical density at 600 nm was measured after 24 hours of growth (for DC and BSM) or 48 hours of growth for (BEP) using a Thermo Genesys 20 spectrophotometer. Cultures containing higher concentrations of crude porcine extract were not saturated in 24 hours. In all cases, cells settled at the bottom of the culture were resuspended before optical density was measured. Cultures containing higher concentrations of BEP were vortexed vigorously before measuring density to disrupt biofilm aggregates that developed during growth. To monitor growth kinetics, 200 ul of the 3 ml cultures described above were transferred to a 96-well plate and incubated in a Tecan Infinite M Nano plate reader housed in an anaerobic chamber. Absorbance was measurements every 10 minutes during growth. Bile salt mixture and deoxycholate stock solutions (5% w/v) were prepared in water and filter sterilized. Porcine bile extract solutions were also prepared in water (5% w/v) as follows. After shaking overnight at 37°C to facilitate solubilization, the solution was centrifuged to clear insoluble material, sterilized by filtration and stored in the dark. Before use, the filtered BEP was incubated at 37C for at least 1 hour to resolubilize components that separate from solution at room temperature.

### B. fragilis P207 genome sequencing

The complete *B. fragilis* strain P207 genome was produced by first combining published P207 metagenome sequence (24) with long-read sequences collected from an Oxford Nanopore MinIon device. Assembly of long reads plus metagenome sequence was carried out using Flye v2.8 (39) with the metagenome option. Paired-end short reads from an Illumina HiSeq 4000 were then used to polish the Flye assembled genome using Pilon v1.23 (40). This assembly approach yielded a circular genome of 5,040,211 base pairs. The complete genome sequence is available through NCBI GenBank accession CP114371. Reads used to assemble the genome are available at the NCBI Sequence Read Archive (accessions SRR22689962 & SRR22689963.

### Transcriptomic analysis of the B. fragilis P207 bile response

An overnight starter cultures of *B. fragilis* P207 was grown from a freezer stock in BHIS at 37°C in a Coy anaerobic chamber. The starter cultures were diluted into triplicate tube with 20 ml fresh BHIS to a starting OD_600_ of 0.05 and outgrown for 6 hours to an OD_600_ of approximately 0.3. Three ml of each culture was harvested for the untreated control, then 1 ml of 0.17% DC in BHIS was added to yield a final concentration of 0.01%. Six minutes after mixing, 3 ml culture was harvested. Again 20 minutes after mixing 3 ml of culture was harvested. Harvesting entailed immediate removal from the anaerobic chamber, centrifugation at 15,000 x g for 1 minute, removal of supernatant and immediate resuspension of the cell pellet in 1 ml TRIzol. Samples were then stored at -80°C until RNA extraction. RNA purification entailed heating the TRIzol samples at 65°C for 10 min followed by addition of 200 μl chloroform, vortexing and incubation at room temperature for 5 min. Aqueous and organic phases were separated by centrifugation at 17,000 x g for 15 min. The upper aqueous phase was transferred to a fresh tube. 0.7X volumes of 100% isopropanol were added, and samples were stored at -80°C overnight. Samples were then centrifuged at 17,000 x g for 30 min at 4°C to pellet the nucleic acid. Pellets were washed twice with 70% ethanol and allowed to air dry before resuspension in 100 μl RNase-free water. DNAse treatment, stranded library preparation using Illumina’s Stranded Total RNA Prep Ligation with Ribo-Zero Plus kit and custom rRNA depletion probes, and sequencing (2 X 50 bp paired end reads using NextSeq2000) was performed by the Microbial Genome Sequencing Center (Pittsburgh, PA). Reads were mapped to the *B. fragilis* P207 genome (GenBank accession number CP114371) using CLC Genomics Workbench 22 (Qiagen). Differential gene expression False Discovery Rate (FDR) p-values were calculated using the method of Benjamini and Hochberg (91). A cutoff criterial used to identify transcriptionally regulated genes was an absolute transcript log_2_(fold change) > 1.5 and FDR p-value < 10^−10^. RNA sequencing reads are available NCBI GEO GSE220692.

We performed RT-qPCR on select genes using the Luna Universal One-Step Kit (New England Biolabs) and QuantStudio 5 (Thermo). Gene specific primers are listed in Table S7. The RNA used in the RNAseq analysis was used as the template and each reaction was run in technical triplicate. The RT-qPCR reaction conditions were as follows: 55°C for 10 min, 95°C for 1 min, 40 X (95°C for 10 s, 60°C for 30 s), followed by a thermal melt to verify product purity. The threshold cycle (Ct) values for each reaction were determined using QuantStudio software (Thermo). The Ct of a sigma-70 RNA polymerase sigma factor (PTOS_001202) was used as the reference for normalization. Expression changes (-ΔΔCt) were calculated using the following relationship: - ((Ct_x_-Ct_ref_)_20 min_-(Ct_x_-Ct_ref_)_0 min_).

### Pathway tools analysis of RNAseq data: assignment of Interpro and GO terms and pathway enrichment analyses

Interpro (49) and GO terms (50) were assigned to each gene in the *B. fragilis* P207 genome using BioBam Cloud BLAST (Table S2). We then implemented both gene set enrichment analysis (GSEA) (51), and Fisher’s exact test (52) to identify gene function classes that are significantly enriched in up- or down-regulated genes after 20 minutes of DC treatment.

### Construction of a pooled barcoded B. fragilis P207 Tn-Himar mutant library

Barcoded himar transposons were introduced into *B. fragilis* P207 from the *E. coli* donor strain, AMD776, carrying the pTGG46-NN1 himar transposon vector library (53) (gift from Adam Deutschbauer, University of California-Berkeley, USA) via conjugation. Briefly, a 10-12 ml BHIS starter culture of *B. fragilis* P207 was inoculated directly from a freezer stock and grown anaerobically for ∼16 hours. Simultaneously, 6 ml LB-C-DAP was inoculated with 0.3 ml of a thawed AMD776 pool and shaken for ∼16 hours. The *B. fragilis* starter culture was diluted into 150 ml fresh BHIS containing 20 ug/ml gentamicin and grown anaerobically for 6-8 hours while 0.5 ml of the *E. coli* starter was diluted into 100-150 ml of fresh LB-C-DAP and grown aerobically for the same duration. The optical density of each culture was measured at 600nm and the cultures were mixed at approximately 1:1 normalizing by OD. Cells were collected by centrifugation (7,000 g for 4 min) and resuspended in a total of approximately 5 ml fresh BHIS. This cell slurry was transferred in 60-70 spots of 100 ul each to 10-12 BHIS plates containing 0.3 mM diaminopimelic acid to support growth of the *E. coli* donor strain. When the liquid absorbed, the plates were incubated aerobically for ∼12 hours and then transferred to the anaerobic incubator for ∼12 hours. Groups of 10-12 mating spots were scraped from the plates, resuspended together in 3 ml BHIS and spread evenly over 10-12 BHIS plates containing erythromycin and gentamicin to select for *B. fragilis* harboring the transposon and aid in counter selection of the donor *E. coli* strain. Plates were incubated anaerobically for 2-3 days. Efficiency of transformation was low; approximately 1 X 10^−7^ *B. fragilis* cells became erythromycin resistant and sets of 10-12 conjugation spots would yield 500-5000 transconjugants. Cells from the sets of 10-12 selection plates in each set were scraped and resuspended in 500 ul BHIS then diluted 10-fold to ensure no cell clumps remained. Then a 450 ul aliquot was inoculated into 45 ml BHIS containing erythromycin and gentamicin and outgrown anaerobically for 10-12 hours. Cells in each outgrowth were collected by centrifugation (7000 g for 4 min), resuspended in 20 ml BHIS containing 30% glycerol and stored in aliquots at -80°C. We eventually obtained approximately 70,000 *B. fragilis* P207 strains harboring Tn-himar insertions in 27 small pools. To combine the mutants in each of the smaller pools, each pool was outgrown in parallel and combined proportional to the approximate number of strains in each pool. Specifically, for each pool, 0.3 ml of thawed glycerol stock was inoculated into 13 ml BHIS containing erythromycin and gentamicin and grown anaerobically for 16 hours. After proportional volumes of each pool were mixed together, glycerol was added to a final concentration of 25%. The mixed pool was distributed in 1 ml aliquots and stored at -80°C. Two aliquots were saved for genomic DNA extraction to map transposon insertion sites.

### Mapping Tn-Himar insertion sites in the B. fragilis BarSeq library

Tn insertions sites were mapped following the approach outlined by Wetmore et al. (54) with modifications to the PCR enrichment. Briefly genomic DNA was extracted using guanidium thiocyanate as previously described (92). Four micrograms of genomic DNA in 130 ul volume were sheared using a Covaris M220 ultrasonicator using manufacturers settings to generate ∼300 bp fragments. Using the NEBNext Ultra II Library Prep Kit (New England Biolabs, E7103S) following the manufacturers protocol, 1 ug of sheared DNA was end-repaired and A-tailed then ligated to a custom Y adapter, prepared by annealing Mod2_TruSeq and Mod2_TS_Univ (Table S7). The final product was cleaned with SPRIselect beads using a two-sided (0.6-0.25X) selection and eluted from the beads in 25 ul 10 mM Tris, pH 8.5. The fragments containing *Himar* transposons were enriched with a two-step nested PCR strategy using Q5 DNA polymerase with GC enhancer (New England Biolabs, M0491) using primers (Table S7) based on primer sequences originally described by Wetmore et al. (54). In the first step, the forward primer contained the Illumina TruSeq Read 1 region, a random hexamer to facilitate clustering, and a transposon-specific sequence that was extended from the original design to improve specificity, and the reverse primer contained the Illumina TruSeq Read 2 sequence complementary to the adapter. In the second step, the primers result in the addition of Illumina P5 and P7 sequences as well as a 6-bp index on the P7 end. Reaction 1 contained 0.5 uM of each of the primers (TS_bs_T7-35 and TS_R), 0.2 mM dNTP, 1X Q5 reaction buffer, 1X GC enhancer, 2 Units Q5 polymerase and 10 ul of adapter-ligated fragments in a 100 ul reaction volume. Cycling parameters were 98°C for 3 min, 20 X (98°C 30 s, 66°C 20 s, 72°C 20 s), 72°C for 5 min, 4°C hold. The first reaction was cleaned with 0.9X AMPure XP beads (Beckman Coulter, A63880) and eluted in 30 ul of 10 mM Tris, pH 8.5. Reaction 2 contained 0.5 uM of each primer (P5_TS_F and P7_MOD_TS_index6), 0.2 mM dNTP, 1X Q5 reaction buffer, 1X GC enhancer, 2 Units Q5 polymerase and 15 ul of cleaned reaction 1 in a 100 ul reaction volume. Cycling parameters were 98°C for 3 min, 15 X (98°C 30 s, 69°C 20 s, 72°C 20 s), 72°C for 5 min, 4°C hold. The final product was cleaned with SPRIselect beads using a two-sided (0.9-0.5X) selection and eluted from the beads in 40 ul 10 mM Tris, pH 8.5. The amplified fragments were sequenced using a 150-cycle MiniSeq High Output Reagent Kit (Illumina, FC-420-1002) on an Illumina MiniSeq. To aid in clustering, sequencing runs were supplemented with 20-30% phiX DNA. Sequences were analyzed using custom scripts written and described by Wetmore and colleagues (54) and available at https://bitbucket.org/berkeleylab/feba/src/master/.

Briefly, the locations of Himar transposon insertions were aligned and mapped to the *B. fragilis* P207 chromosome sequence (NCBI accession CP114371) using BLAT, and unique barcode sequences were associated with their corresponding genome insertion location using the custom Perl script, MapTnSeq.pl. Sets of barcodes that reliably map to one location in the genome were identified using the custom Perl script, DesignRandomPool.pl with the following flags -minN 8 -minFrac 0.7 -minRatio 7. Mapping statistics are provided in Table S3. Raw Tn-seq data are deposited in the NCBI sequence read archive under BioProject accession PRJNA910954; BioSample accession SAMN32154224; SRA accession SRR22677646.

### Assessment of B. fragilis P207 gene essentiality in BHIS medium

The *B. fragilis* P207 Tn-himar library was prepared using BHIS growth medium. Quantifying the frequency of insertions across the genome can provide some indication of gene essentiality when cells are grown on BHIS. Analysis of essentiality was performed using the TRANSIT package (available at https://github.com/mad-lab/transit) (93). Briefly, counts of transposon insertions at individual TA dinucleotides sites were measured with the TPP tool in TRANSIT, which uses the BWA aligner (94) to map *himar-B. fragilis* junctional reads to the *B. fragilis* P207 genome. Based on these count data, gene essentiality was calculated using both Gumbel (56) and HMM (55) methods in the TRANSIT package. Output from this analysis is available in Table S5.

### Construction of an arrayed B. fragilis P207 Tn-Himar mutant library

We assembled a limited collection of individual *B. fragilis* P207 *Tn-Himar* mutant strains arrayed in 96-well plates. The mutants were generated by conjugating barcoded transposons from AMD776 into *B. fragilis* P207 as described above. However, instead of pooling transconjugants, individual colonies were manually picked into deep well plates containing 1 ml BHIS with 5 ug/ml erythromycin per well. Plates were incubated anaerobically overnight until wells became turbid. In a fresh deep well plate, 0.5 ml each culture was mixed with 0.5 ml sterile 50% glycerol to yield a final concentration 25% glycerol. The plate with glycerol was sealed with adhesive foil seals and stored at -80°C. A small aliquot from the initial culture plate was saved at - 20°C for mapping the transposon insertion sites in each clone (see below). A total of 1020 clones were picked from 2 independent conjugations. This yielded a collection of 889 successfully mapped *B. fragilis* P207::himar clones representing 723 unique insertion sites. Insertions in 824 clones mapped to predicted coding regions (670 unique sites in 575 unique genes). Thus approximately 14% of genes are represented in this collection. The remaining 65 clones had insertions in predicted non-coding regions. These mapped arrayed mutants are cataloged in Table S4.

### Mapping insertion sites in individual B. fragilis P207 Tn-Himar mutants

Insertion sites in individual clones were mapped using a two-step nested arbitrary PCR approach using 2X GoTaq master mix (Promega M7122). The first reaction (1X GoTaq master mix, 0.3 uM U1 fw, 0.3 uM M13-N7 and 0.5 ul saved culture in a 20 ul reaction) was cycled using the following parameters: 95°C for 2min, 35X (95°C for 30 sec, 38°C for 30 sec, 72°C for 1 min), 72°C for 5 min, 12°C hold. The products were enzymatically cleaned (3 ul of PCR reaction, 2 ul water, 1 ul of ExoSap-IT (Applied Biosystems 78200)) for 15 minutes at 37°C followed by 15 minutes at 80°C. The second PCR reaction (1X GoTaq master mix, 0.3 uM U2 out, 0.3 uM M13Fw and 2 ul of cleaned product above in a 20 ul reaction) was cycled using the following parameters: 95°C for 2 min, 35X (95°C for 30 sec, 68-48°C dropping 0.5°C per cycle for 30 sec, 72°C for 1 min), 72°C for 5 min, 12°C hold. Secondary PCR products were treated with ExoSap-IT as above and Sanger sequenced using the U2 out primer. Sequences were compared to the genome using BLAST to identify insertion sites.

### Cultivation of the Tn-Himar library in bile

A 0.5 ml aliquot of the pooled *B. fragilis* mutant library glycerol stock was inoculated into 25 ml BHIS containing 5 ug/ml Erythromycin and 20 ug/ml Gentamycin and outgrown anaerobically overnight. Cells from four aliquots of 1 ml each were collected by centrifugation (2 min at 12,000 g), resuspended in 1 ml of phosphate-buffered saline to wash residual media, centrifuged again (2 min at 12,000 g) and stored in pellets -20°C as reference samples. Then 20 ul of overnight culture (containing approximately 2 X 10^8^ CFU) were inoculated into quadruplicate tubes containing 4 mL of either BHIS, BHIS with 0.01% bile salt mix, BHIS with 0.01% deoxycholate. These concentrations reduced growth of wild-type cultures by approximately 20% or 30% respectively (Figure S1). Cultures were incubated anaerobically for 24 h, which allowed for approximately 7 doublings. We harvested cells from 1 ml of these cultures, washing the cell pellets with PBS as we did for the reference samples. In addition, to amplify fitness differences between strains in the mutant pool, we back-diluted these cultures into the same conditions (100 ul into 4 ml fresh media) and allowed a second 24 hour growth period to allow further differentiation of the fitness of individual mutant strains. Again, 1 ml of the passaged day 2 cultures were harvested as above. To assess strain fitness we amplified and sequenced the transposon barcodes after the first and second passages of the pool in the untreated and treatment conditions (see below).

We conducted a similar experiment using porcine bile extract. However, because we experienced more variability in growth with porcine bile extract, we conducted this experiment with three different concentrations, 0.04, 0.08 and 0.16%, using a similar passaging approach from an independent outgrowth of the library. To ensure a maximal number of cell doublings, cultures were incubated until saturation, which was longer than 24 hours at the higher concentrations of porcine bile extract. For the porcine bile extract treatments, barcode abundance was assessed after the second passage.

### Amplification and sequencing of Tn-Himar barcodes

To assess barcode abundances, we followed the approach developed and described by Wetmore and colleagues (54). Briefly, each cell pellet was resuspended in approximately 50 ul water. Barcodes were amplified using Q5 polymerase (New England Biolabs) in 20 ul reaction volumes containing 1X Q5 reaction buffer, 1X GC enhancer, 0.8 U Q5 polymerase, 0.2 mM dNTP, 0.5 uM of each primer and 1 ul of resuspended cells. Each reaction contained the Barseq_P1 forward primer and a unique indexed reverse primer, Barseq_P2_ITxxx, where the xxx identifies the index number (54). Reactions were cycled as follows: 98 °C for 4 min, 25X (98 °C for 30 s, 55 °C for 30 s, and 72 °C for 30), 72 °C for 5 min, 4°C hold. PCR products separated on a 2% agarose gel to confirm amplification of a 190 bp product. Aliquots (5 ul) of each reaction were pooled. The pool of PCR products was centrifuged for 2 min at 16,000g to pellet cell debris. Then 90 ul of the mix was transferred to a fresh tube and cleaned with SPRIselect beads (Beckman Coulter) using a two-sided (1.2x-0.5X) selection. The amplified barcodes were sequenced using a 75-cycle MiniSeq High Output Reagent Kit (Illumina, FC-420-1001) on an Illumina MiniSeq at Michigan State University. To aid in clustering, sequencing runs were supplemented with 20-30% phiX DNA. Sequence data have been deposited in the NCBI Sequence Read Archive under BioProject accession PRJNA910954; BioSample accession SAMN32154224; SRA accession SRR22686218-SRR22686272.

### Analysis of Tn-Himar strain fitness

Barcode sequences were analyzed using the fitness calculation protocol of Wetmore and colleagues (54). Briefly, the barcodes in each sample were counted and assembled using MultiCodes.pl and combineBarSeq.pl. Then using the barcode abundance data and the mapping information for the mutant pool, gene fitness was calculated using the R script, FEBA.R. The fitness of each strain was calculated as a normalized log_2_ ratio of barcode counts in the treatment sample to counts in the reference sample. The fitness of genes was calculated as the weighted average of strain fitness values, the weight being inversely proportional to a variance metric based on the total number of reads for each strain; this approach is fully described in (54). Insertions in the first 10% or last 10% of a gene were not considered in gene fitness calculations. The complete data set of fitness values and t scores for each condition is listed in Table S6.

To identify genes that contribute to fitness in any bile condition, we filtered the fitness scores in each condition to include scores greater than 1.5 or less than -4 after the second passage. We then manually removed genes which resulted in fitness defects in the absence of bile (average fitness in BHIS < -3). Then 10 genes for which the fitness defect with bile is not substantially worse than without bile (fitness in bile minus fitness defect without bile > -3) were manually removed (PTOS_000072, PTOS_000680, PTOS_001138, PTOS_002708, PTOS_002806, PTOS_003551, PTOS_003802, PTOS_003809, PTOS_003811, PTOS_004091). This resulted in 122 unique genes whose fitness scores were hierarchically clustered using uncentered correlation and average linkage using Cluster 3.0. Heatmap of clustered fitness scores was rendered with Prism (GraphPad) with some manual rearrangement of genes to bring together genes that are adjacent on the chromosome and have similar fitness profiles.

## Acknowledgements

This work was supported by an RC2 award (5RC2DK122394) from the National Institute of Diabetes and Digestive and Kidney Diseases (NIDDK) to EC, and a MIRA award (R35GM131762) to SC from the National Institute of General Medical Sciences. The University of Chicago Digestive Disease Research Core Center is supported by a P30 award (P30DK042086) from the NIDDK. The funders had no role in study design, data collection and interpretation, or the decision to submit the work for publication.

## Supplemental Materials

**In separate documents:**

**Table S1: RNAseq expression data.xlsx**

**Table S2: Overrepresented_functional_groups_in_up_and_downregulated_genes.xlsx**

**Table S3: Mapping statistics**

**Table S4: P207-TN_arrayed_collection.xlsx**

**Table S5: Essential gene calls.xlsx**

**Table S6: BarSeq output.xlsx**

**Table S7: Strains and primers**

### Supplemental Results

#### Multiple classes of transcription factors are regulated by DC

Approximately one quarter of all *B. fragilis* P207 genes exhibited significant differential expression upon acute DC exposure. The massive transcriptional response that we observe upon DC treatment is likely a result of a cascade of transcriptional regulatory events cued by stress inflicted by deoxycholate. Two of the top three enriched functional categories in the activated gene set relate to transcription (enriched GO terms: transcription cis-regulatory region binding, and regulation DNA-templated transcription; Table S2). The *B. fragilis* sigma factors had a striking profile in which eight sigma factors were activated and ten repressed (|log_2_(FC)|>1.5) by the 20 minute time point, (Figure S6A; Tables S1 & S2). Other common classes of transcriptional regulators included in the enriched gene set include one-component (1) and two-component system (2) genes, which show a similarly disparate regulatory profile as the alternative sigma factors (Figure S6 B-C; Tables S1 & S2), indicating a complex response to DC exposure.

#### Transcriptional evidence for a metabolic shift upon DC exposure

By far, the most significantly enriched functional categories in the down-regulated set of genes involve translation (enriched GO terms: ribosome, translational elongation, aminoacyl-tRNA ligase activity, tRNA aminoacylation for protein translation). Expression of ribosomal proteins, ribosomal accessory factors, and aminoacyl tRNA synthetases is uniformly lower (by 2-50 fold), with few exceptions (main text Figure 1B-C, Tables S1 & S2). We note that most of these genes are not represented in our fitness data as translation is an essential process. The transcriptional effect observed resembles gene expression changes induced by the stringent response (3), which slows growth and facilitates cellular adaptation to nutrient or energy limitation. The wholesale downregulation of protein expression machinery is accompanied by reduced expression of genes required for cell growth and division (enriched GO terms: regulation of cell shape, cell wall organization, peptidoglycan biosynthetic process, lipopolysaccharide biosynthetic process, cell envelope) (Tables S1 & S2). Other significantly down-regulated biosynthetic pathways (FDR q-value < 0.05) include genes required for production of cobalamin, ribonucleosides, pseudouridine, pantothenic acid, queuosine, arginine, and coenzyme A.

While the transcriptomic data provide evidence that deoxycholate treatment results in a large-scale downregulation of anabolic metabolism, expression of select metabolic enzymes is activated. Among the most notable of these responses is the transcriptional activation of genes involved in glutamate metabolism (enriched GO term: glutamate metabolic process) (Table S2). Specifically, DC treatment induces expression of asparagine synthase B (*asnB; ptos_002403*), glutaminase (*glsA; ptos_000368*), genes in the histidine utilization pathway (*ptos_003734-3738*), and glutamate dehydrogenase (*gdhA; ptos_003163*), all of which catalyze production of glutamate from various substrates.

The expression of glutamate decarboxylase (GAD, *ptos_000367*), which converts glutamate to γ-amino butyric acid (GABA), is also activated. This reaction consumes a proton and is thought to contribute to neutralization of an acidified cytoplasm (4). Glutamate decarboxylase is associated with tolerance of acid and bile acid stress in many bacteria, including *Bacteroides* species (5-7). Most of these genes have neutral fitness scores. However strains with disruptions in *gdhA*, whose activity results in the generation of reducing equivalents, have a fitness advantage. This result is consistent with the fitness advantage of disrupting other pathways that result in the generation of reducing equivalents, namely the oxidative phase of the pentose phosphate pathway (*zwf, pgl, gnd*), and the first step of beta-oxidation of fatty acids (*fadE* and *fixAB*) (main text Figure 3 & 5).

A second set of upregulated metabolic genes have predicted roles in citrate/carboxylate metabolism (enriched GO term: tricarboxylic acid cycle) including the gene cluster *ptos_003272-3274* (citrate synthase, isocitrate dehydrogenase, and aconitase) and a succinyl-CoA ligase system (*ptos_001896-1897*). Activation of lactate/malate dehydrogenase (*ptos_000443*) and L-lactate permease (*ptos_001319*) expression by approximately 50-fold in the presence of DC indicates a shift in carboxylic acid metabolism in *B. fragilis* during bile stress. Finally, transcription of multiple enzymes comprising the pentose phosphate pathway (enriched GO term: pentose-phosphate shunt) including *gnd, zwf*, and *pgl* (*ptos_1536-1538*), *fba* (*ptos_002818*) and *rpiA* (*ptos_003205*) are activated by DC. Other genes in this pathway including *rpiB* (*ptos_001356*) and *ptos_002671* and *ptos_002867* are repressed by DC (Table S1). This pathway produces key substrates for anabolic metabolism and also yields reducing equivalents.

#### DC-regulated expression of membrane ion transport systems

Transcription of the F_O_F_1_ ATP synthase operon (*ptos_001821-1829*) is activated upon DC exposure (enriched GO terms: proton motive force-driven ATP synthesis), suggesting that DC compromises energy production in *B. fragilis*, perhaps by affecting membrane permeability. Indeed, expression of multiple genes with predicted roles in gated ion transport (enriched GO terms: monoatomic ion gated channel activity) (Table S2) is activated by DC including ion channels *ptos_003102* and *ptos_000442*, mechanosensitive ion channels *ptos_002809, ptos_004068, ptos_000516, ptos_000659, ptos_000815*, and the H(+)/Cl(-) exchange transporter *ptos_001050*. Overall, we observe a variable pattern of activation and repression of annotated ion transporters with the highest level of activation (≈20-fold) for a Na+/H+-translocating pyrophosphatase family protein, *ptos_000210* (8, 9), and transporters with predicted roles in sulfate (*ptos_002300*) and zinc transport (*zup; ptos_002150*). The large effect of DC treatment on ion transport gene transcription provides evidence of general dysregulation of cellular ion homeostasis.

#### B. fragilis P207 relies on multiple efflux and stress response systems in the presence of bile

Efflux is a well-established mechanism of bile-tolerance (10). TolC-family outer membrane proteins work in conjunction with RND- and ABC-family transport systems to enable efflux of a broad range of substrates, including bile acids (11, 12). These tripartite efflux systems are comprised of an inner membrane permease, a TolC-family protein in the outer membrane, and a periplasmic adapter protein that bridges the inner and outer membrane proteins. *B. fragilis* P207 encodes 23 TolC-family exporter systems, most of which are not strongly expressed in BHIS (Figure S8). Previous work in *B. thetaiotaomicon* identified a TolC-type bile-induced efflux system, BT2792-BT2795, that specifically contributed to fitness in the presence of multiple bile acids (13). *B. fragilis* P207 does not encode an orthologous system, but we identified multiple tripartite TolC-containing efflux systems that contribute to fitness in DC and BEP, likely in a redundant fashion (Figure S7). The overall repertoire of tripartite efflux genes that support growth/survival in DC and BEP is similar, though reduced fitness due to disruption of *ptos_000132-34* is specific to DC.

While both our transcriptomic and functional genomic datasets provide evidence that efflux systems are important in the context of bile, these approaches highlight distinct efflux systems (Figure S7). For example, the TolC-family protein encoded by *ptos_003611* is the most highly induced gene in the DC transcriptomic data set (main text Figure 2, Table S1) yet the fitness consequence of disrupting this gene or adjacent transport genes is minor in DC (Figure S7). We note that *ptos_003611-14* does contribute to fitness in crude BEP raising the possibility that DC may act as a signal to upregulate expression of a system that is important for export of other molecules that often co-occur with DC in the mammalian gut. No single system is critical for survival in DC; instead *ptos_000132-34, ptos002878-80* and *ptos_03242-44* each contribute to fitness, likely in a redundant fashion. Transcription of *ptos_003730-32* is activated by DC treatment but insertions in this locus do not impact fitness in our conditions. However, Tn-*Himar* insertions in the adjacent transcriptional repressor gene, *ptos_003733*, confer a fitness advantage in the presence of DC and BEP (Figure S7). From this result, we infer that de-repression of *ptos_003730-32* via disruption of its transcriptional repressor promotes growth and/or survival in DC and BEP. Together these data indicate multiple efflux systems are transcriptionally regulated by and/or contribute to fitness in the presence of bile.

Distinct sets of stress response genes are identified in our transcriptomic and functional genomic experiments. Among the most highly activated genes in the transcriptional dataset are those with known roles in acute stress responses, including genes that mitigate protein misfolding (enriched GO terms: unfolded protein binding; protein folding) (Table S2; main text Figure 2A-B). Genes in this class include the chaperones *groEL, groES, htpG, dnaK, clpB*, and a small Hsp20-family protein (*ptos_002388*). Most of these genes are not represented in our fitness data (Table S6, Figure S9). On the other hand, transcripts corresponding to the protein chaperone *dnaJ* (*ptos_001434*), which is important for maintaining proteostasis during stress, are only modestly elevated in DC, but strains with disruptions in this gene are strongly attenuated in all bile conditions. The ClpXP protease system encoded by *ptos_003701-2* also fosters protein quality control during stress. Similar to *dnaJ*, these genes are critical for fitness in bile, but are not transcriptionally regulated by DC exposure (main text Figure 3, Figure S9, Table S1 & S6). Thus both data sets provide evidence that growth in bile involves mitigation of protein misfolding. We observe transcriptional responses indicative of oxidative stress in DC (enriched GO terms: response to oxidative stress; cellular oxidant detoxification; and cellular homeostasis) (Table S2). Specifically, *msrB*, which repairs oxidatively-damaged methionine residues, *katB* catalase (*ptos_001055*), superoxide dismutase (*ptos_002119*), peroxide stress protein YaaA (*ptos_003649*) and glutathione peroxidase (*ptos_003459*) are all strongly activated by DC exposure, as are DNA starvation/stationary phase protection protein (*dps; ptos_001118*), and universal stress protein (*usp*; *ptos_002049*) (Table S1, main text Figure 2A-B). However, none of these genes exhibit fitness defects when disrupted (Figure S9, Table S6). Functional redundancy in these oxidative stress responses likely protects cells from loss of any single gene in this group. We further observed induction of all three genes involved in production of inorganic polyphosphate (*ptos_002410, ptos_002960, ptos_003236*), which is known to confer general protective effects during stress exposure in a variety of bacterial taxa (14-17). Again, disruption of any one of these three polyphosphate kinases is not detrimental in any condition tested (Figure S9, Table S6). Finally, the constitutively expressed *bat* genes (*ptos_02052-58*), which have a reported role in aerotolerance (18), are critical for fitness in our bile conditions (main text Figure 3, Figure S9, Table S6).

#### Candidate Essential Genes of Note in B. fragilis P207

We note several candidate essential genes that we found to be of particular interest. Transposon insertions were not recovered in one of the two RelA/SpoT paralogs (*ptos_000629*), which regulate levels of the alarmone ppGpp and metabolic adaptation to stress (3). Similarly, in the closely related species *Bacteroides thetaiotaomicron* one of the RelA/SpoT paralogs is also essential (19). Other notable candidate essential genes include a predicted two-gene operon encoding a type III restriction enzyme (*ptos_000974*) and a type III adenine DNA methyltransferase (*ptos_000975*). This gene pair is highly conserved in *B. fragilis* in contrast to most restriction systems, which are often carried on mobile elements and variable among *B. fragilis* isolates. The *rokA* carbohydrate kinase (*ptos_000462*) has been successfully deleted in a derivative of *B. fragilis* strain 638R (20), but is classified as a candidate essential gene in P207 in our growth conditions based on HMM analysis of insertion data. A *upxY* family transcription antiterminator (*ptos_002531*) is classified as essential by HMM. The orthologous gene in *B. fragilis* 638R (BF638R_2798) was also classified as essential in a global Tn-seq study (21). This *upxY* paralog is not adjacent to a corresponding *upxZ* anti-antiterminator, as is typical in *Bacteroides* capsular polysaccharide loci (22). Rather, *ptos_002531* is the first gene in a putative 28 gene operon that encodes the machinery to synthesize the large capsule EPS elaborated on a fraction of cells in a population (23). Within this large predicted operon, insertion in all downstream genes are recovered, though a gene of unknown function (*ptos_002544*) and a glycosyltransferase family 4 gene (*ptos_002546*) are scored as growth defective based on the distribution of insertions recovered (Table S5).

### Supplemental Figures

**Figure S1:**
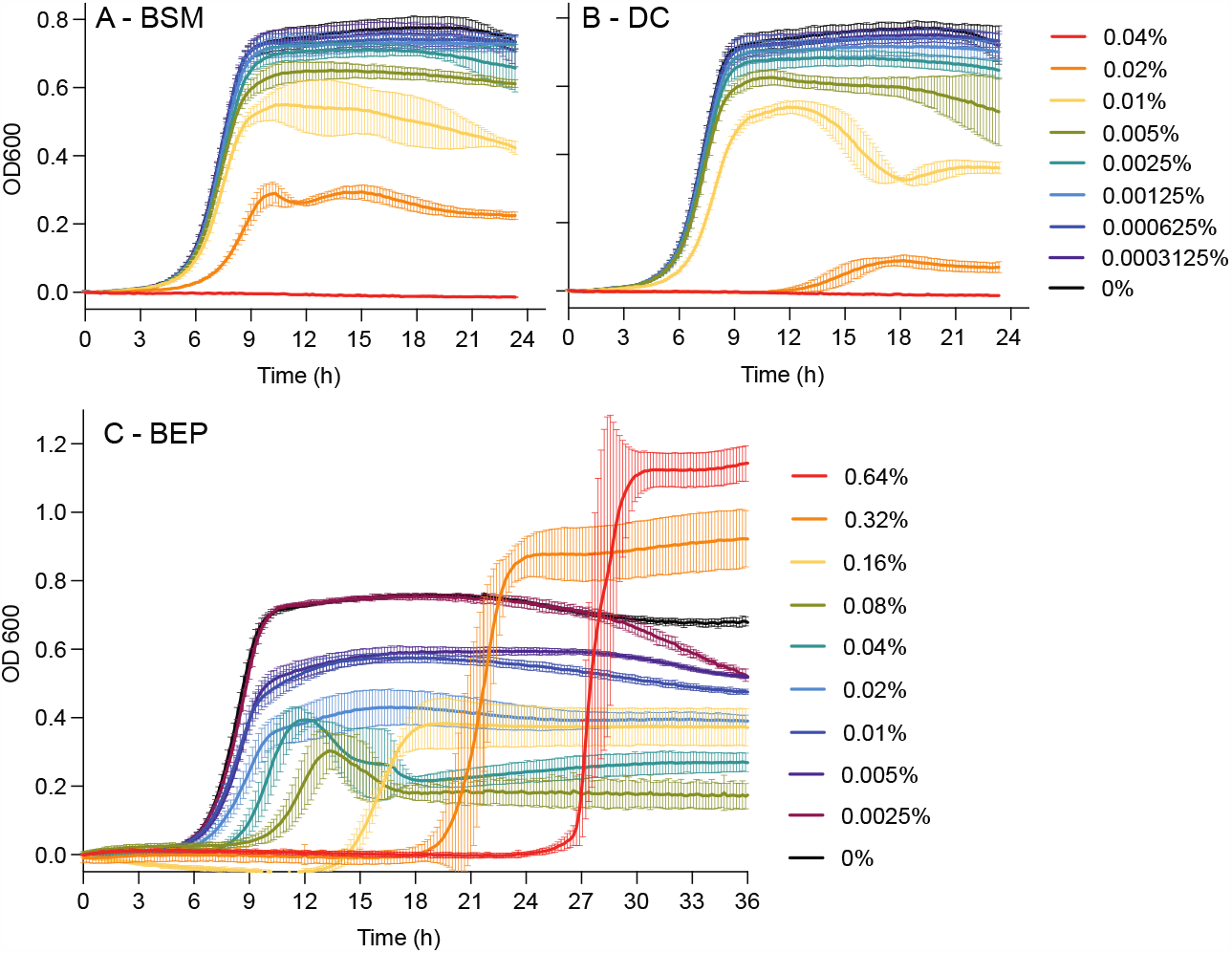
Representative growth curves of *B. fragilis P207* grown in BHIS in the presence of increasing concentrations of bile. **A**. Bile salt mixture (BSM), **B**. deoxycholate (DC), **C**. bile extract of porcine (BEP). Concentrations are in % w/v. The optical density of 200ul of culture in 96-well plates was measured every 10 minutes. Data represent the mean ± SD of 4 replicate cultures grown in the same 96-well plate. The highest concentrations of BEP delayed growth, but enhanced total culture yield.

**Figure S2:**
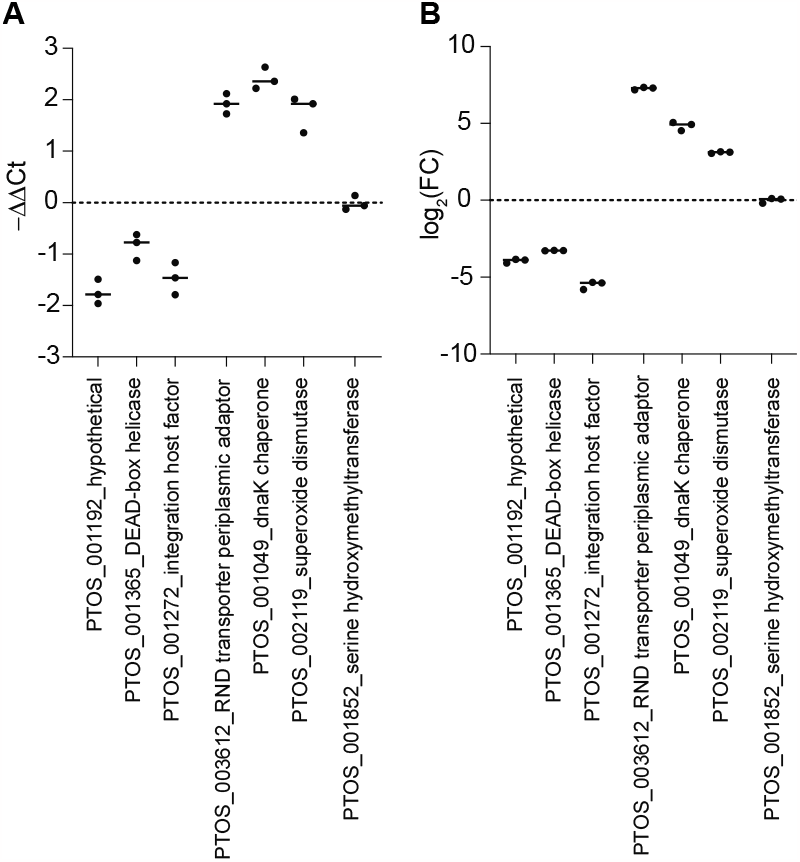
Evaluation of changes in transcript abundance 20 minutes after deoxycholate treatment by RT-qPCR. **A**. Relative abundance of select transcripts evaluated by RT-qPCR compared to a reference gene (PTOS_001202, Sigma^70^ RNA polymerase sigma factor), where -ΔΔCt is the calculated as -((Ct_x_-Ct_ref_)_20 min_-(Ct_x_-Ct_ref_)_0 min_). **B**. Relative abundance of the same set of genes evaluated by RNA seq, where fold change (FC) is average counts CPM_0 min_/average CPM_20 min_. The mean and values of triplicate samples for both approaches are plotted. Ct = threshold cycle. CPM = counts per million counts. For three downregulated, three upregulated and one unchanged gene in or RNAseq dataset, the same directionality of change was observed by RT-qPCR. The magnitude of change detected by these two approaches differs. RNAseq is more sensitive to differences and less variable between replicates.

**Figure S3:**
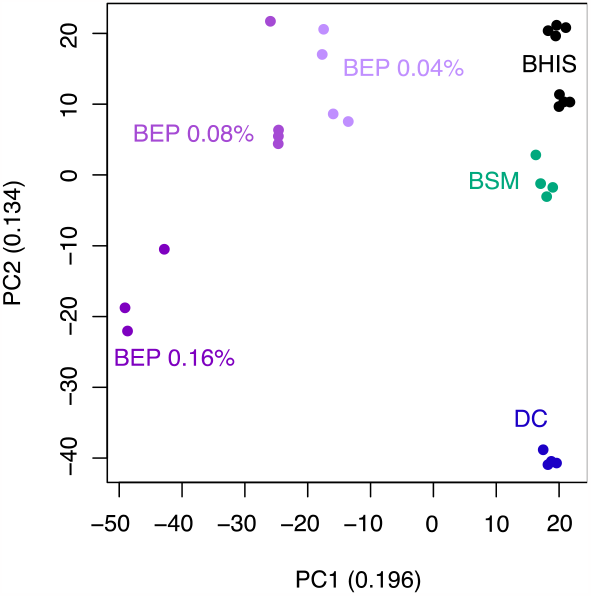
BarSeq replicate data are consistent within each treatment group. Principle component analysis (PCA) of fitness scores from each experimental replicate treatment after the second passage of growth in deoxycholate (DC), bile salt mixture (BSM), plain BHIS, and 0.04%, 0.08%, and 0.16% bile extract porcine (BEP). Genes lacking fitness scores in any experimental set were removed before PCA. The first two principal components are plotted with the fraction of variance accounted for on each axis.

**Figure S4:**
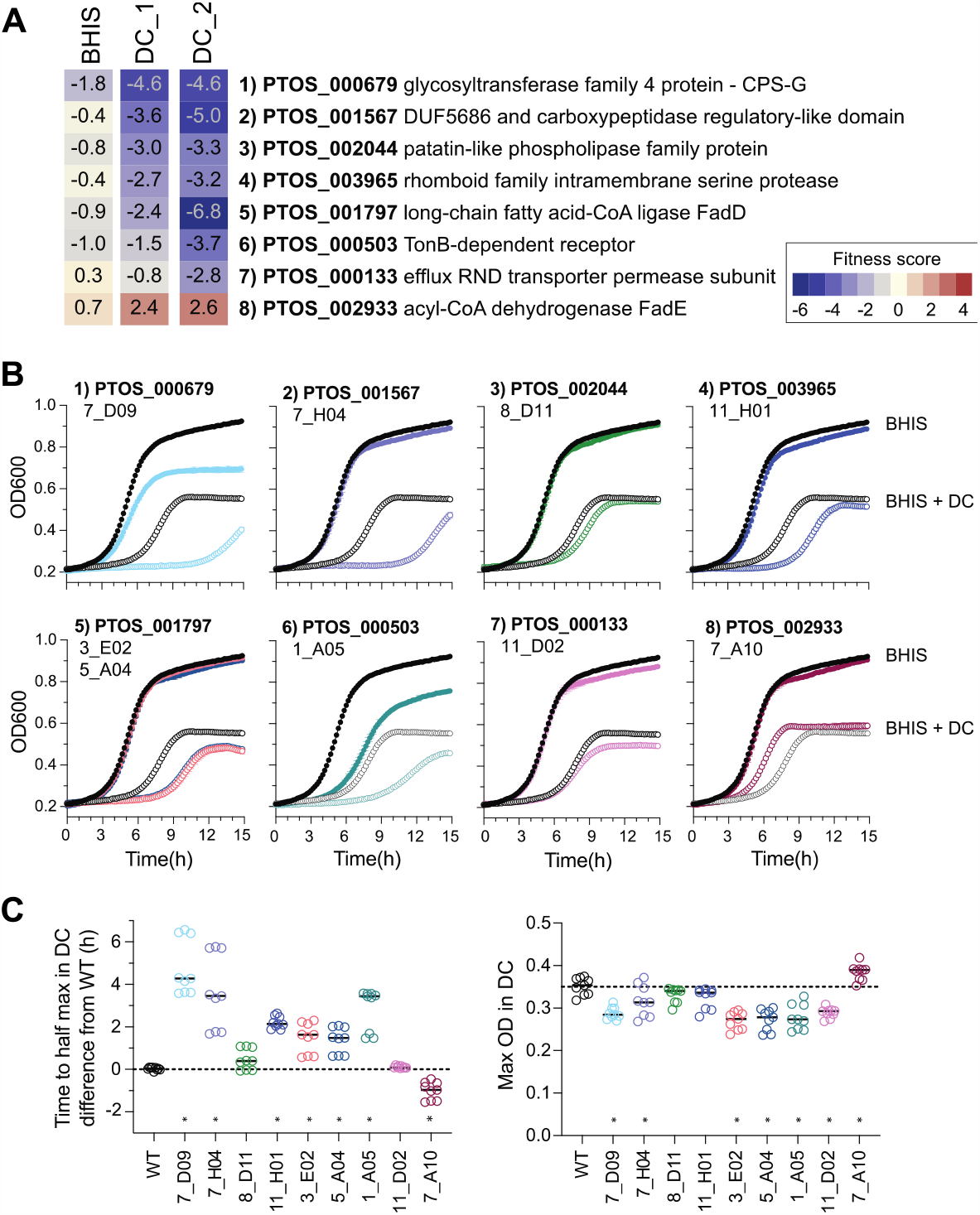
Individual transposon mutant strains validate fitness phenotypes observed in mutant pool. **A**. Heat map of composite BarSeq fitness scores for 8 genes that were represented by clones in our arrayed collection of mapped transposon mutant strains. Fitness scores after first passage in BHIS or BHIS with DC (DC-1), and also after second passage in BHIS containing DC (DC_2) are indicated. Genes were ranked by fitness score after first passage in DC (DC_1), which most similarly reflects the growth curves below. **B**. Representative growth curves of WT (black circles) and mutant clones (colored circles) grown in BHIS without (closed circles) and with (open circles) 0.01% DC. Growth was monitored by absorbance (OD_600_) every 10 minutes and data represent mean ± SD of triplicate cultures assayed together. Error bars are often smaller than, and masked by, the symbol. One of three independent trials is presented. The genes and corresponding mutant clone names are indicated in each graph. Our collection contained two strains with insertions in *ptos_001797*, represented by different colors. Most mutants grow similar to WT in the absence of DC, although strains with insertions in *ptos_000679* or *ptos_000503* exhibit defects in BHIS broth alone, consistent with the negative fitness scores in BHIS. **C**. We calculated growth parameters using the Growthcurvr package in R on the growth curves in BHIS+DC on triplicate cultures from three independent trials (n=9). The difference in time to half-max compared to wild-type (t_mid_mut_ – t_mid_WT_), which reflects a composite of differences in lag and growth rate is shown on the left. The max density (k) in BHIS+DC of each growth is shown on the right. In both cases, difference to WT was assessed by one way ANOVA followed by Dunnett’s multiple comparisons test (* p<0.05). All mutants are statistically different from WT in at least one of these measurements, except 8_D11. The transposon insertion in this strain is in the last 13% of PTOS_002044. It is not uncommon for C-terminal truncations to have modest or no effects. In fact, composite fitness scores ignore strains with insertions in the first or last 10% of a gene because insertions in the termini of gene regions often do not fully disrupt gene function. Overall, the negative fitness scores for genes correspond to either delayed growth and/or lower overall growth in DC for the individual insertion strains compared to WT. Importantly, our collection also contained a strain with an insertion in FadE, which has a positive fitness score in the bulk passaging experiment. The individual strain with an insertion in this gene consistently outperformed WT in the presence of DC, reaching half-max on average an hour sooner than WT and reaching a higher terminal density than WT.

**Figure S5:**
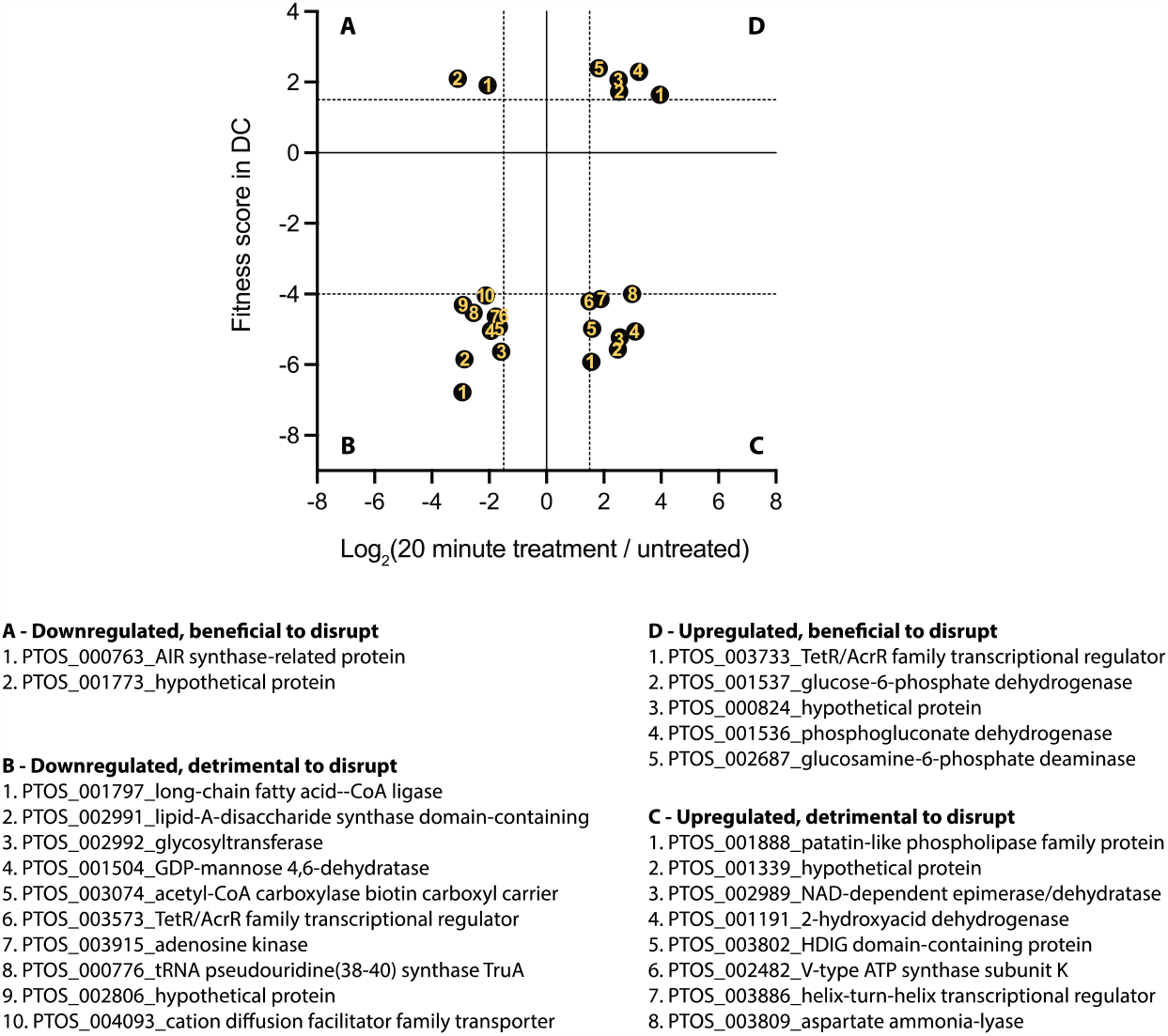
Transcriptionally regulated genes with significant fitness scores. This graph is identical to Figure 4D in the main text, except that the genes in each quadrant (A-D) are numbered. Below the graph are the locus number and predicted annotation corresponding to the numbered genes in each quadrant.

**Figure S6:**
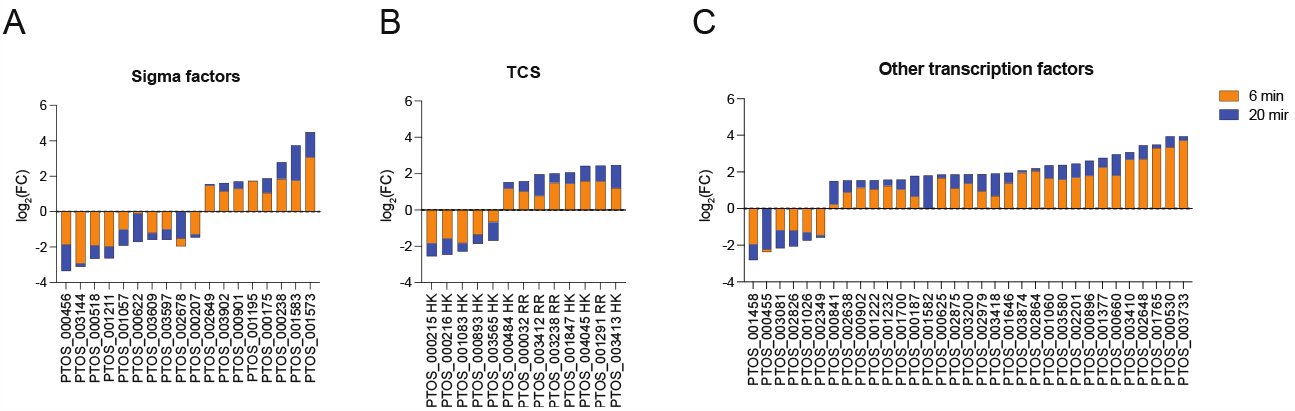
Transcriptional regulators exhibit a complex response to DC. Log_2_(fold change) in transcript levels of genes encoding **(A)** alternative sigma factors, **(B)** two-component signaling (TCS) proteins (HK – histidine kinase, RR – response regulator), and **(C)** other transcription factors that are themselves transcriptional regulated after 6 or 20 minutes of DC exposure (orange and blue, respectively).

**Figure S7:**
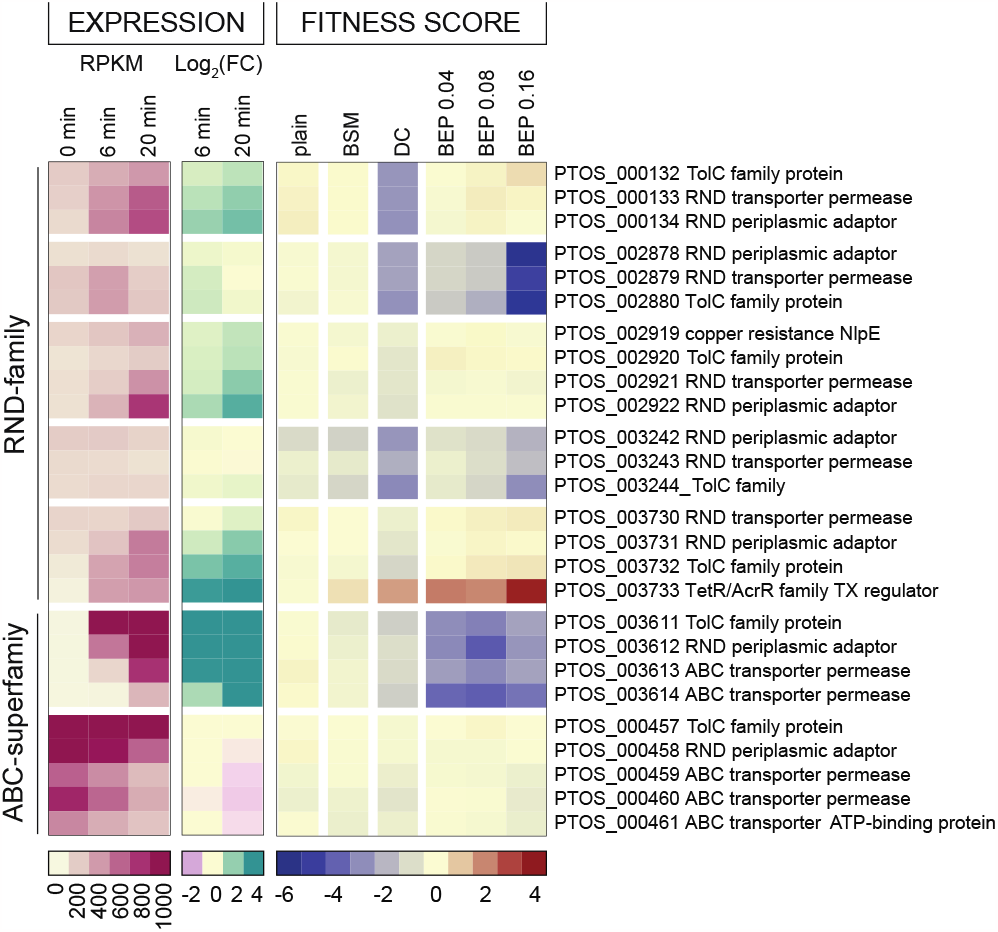
Multiple efflux systems differentially contribute to *B. fragilis* fitness in the presence of purified bile acid and crude bile conditions. Transcriptional and fitness data for TolC-family efflux system that are *either* transcriptionally regulated in the presence of purified deoxycholate *or* contribute to fitness in at least one bile condition (BSM: bile salt mixture, DC: Deoxycholate, BEP: bile extract from porcine). A similar heat map of all 23 TolC-containing operons is presented in Figure S8. Transcript abundance is presented as reads per kilobase per million reads (RPKM) and regulation is indicated by the log_2_(fold change) compared to untreated cells (0 min). Gene-level fitness scores after the second passage in each of the indicated conditions are presented. Color scales are presented below each type of data.

**Figure S8:**
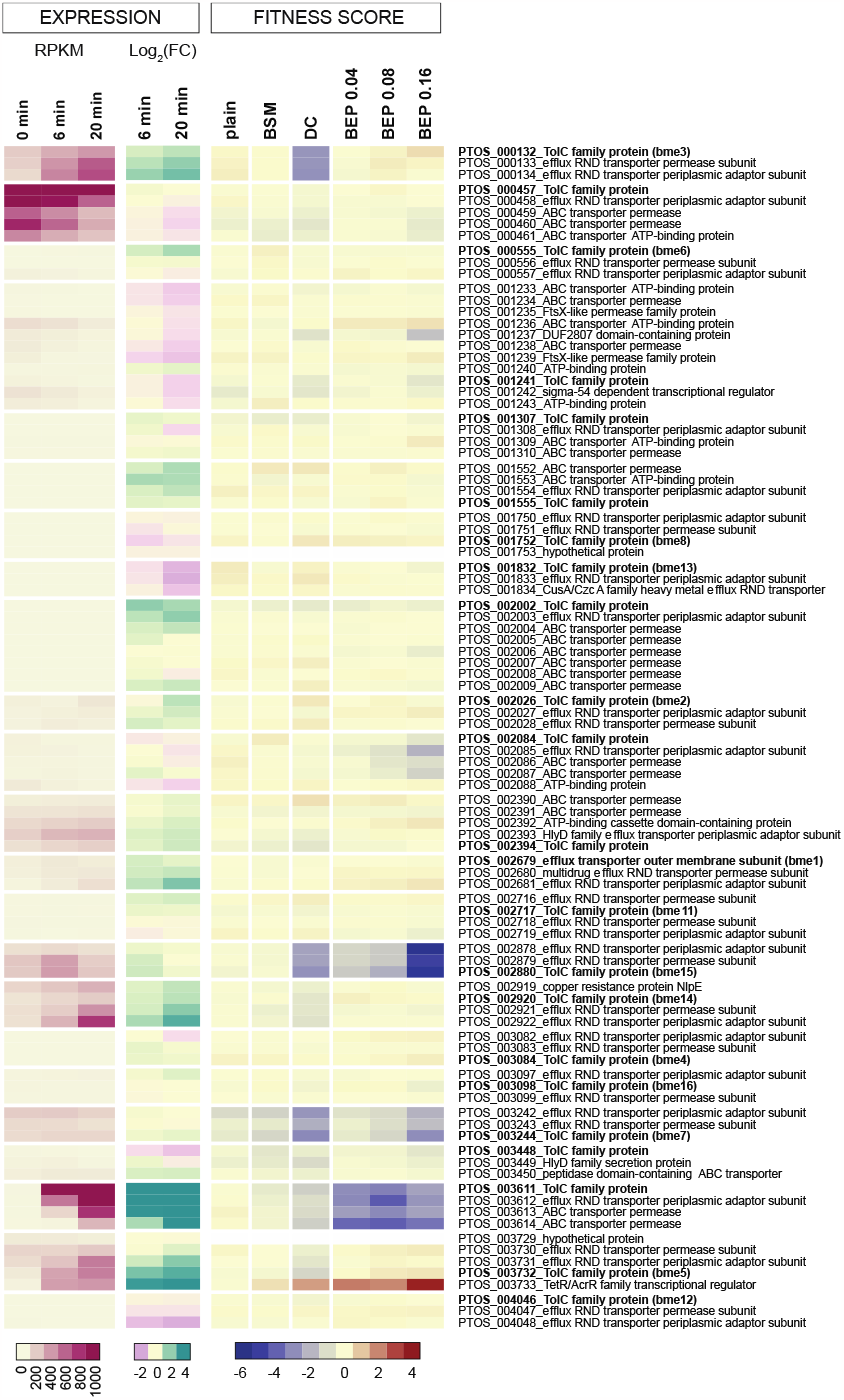
Expression and fitness scores for all 23 TolC-family membrane transport systems. Heat maps transcript levels (RPKM), log_2_(fold change) in transcript and gene-level fitness scores after serial passage in the presence of bile (as in Figure S7) for the operons containing each of the 23 TolC-family exporter genes in the *B. fragilis* P207 genome. TolC-family genes are highlighted in bold. Most tripartite efflux systems are not highly expressed in our *in vitro* conditions. Only a small subset contribute to fitness in our bile conditions.

**Figure S9:**
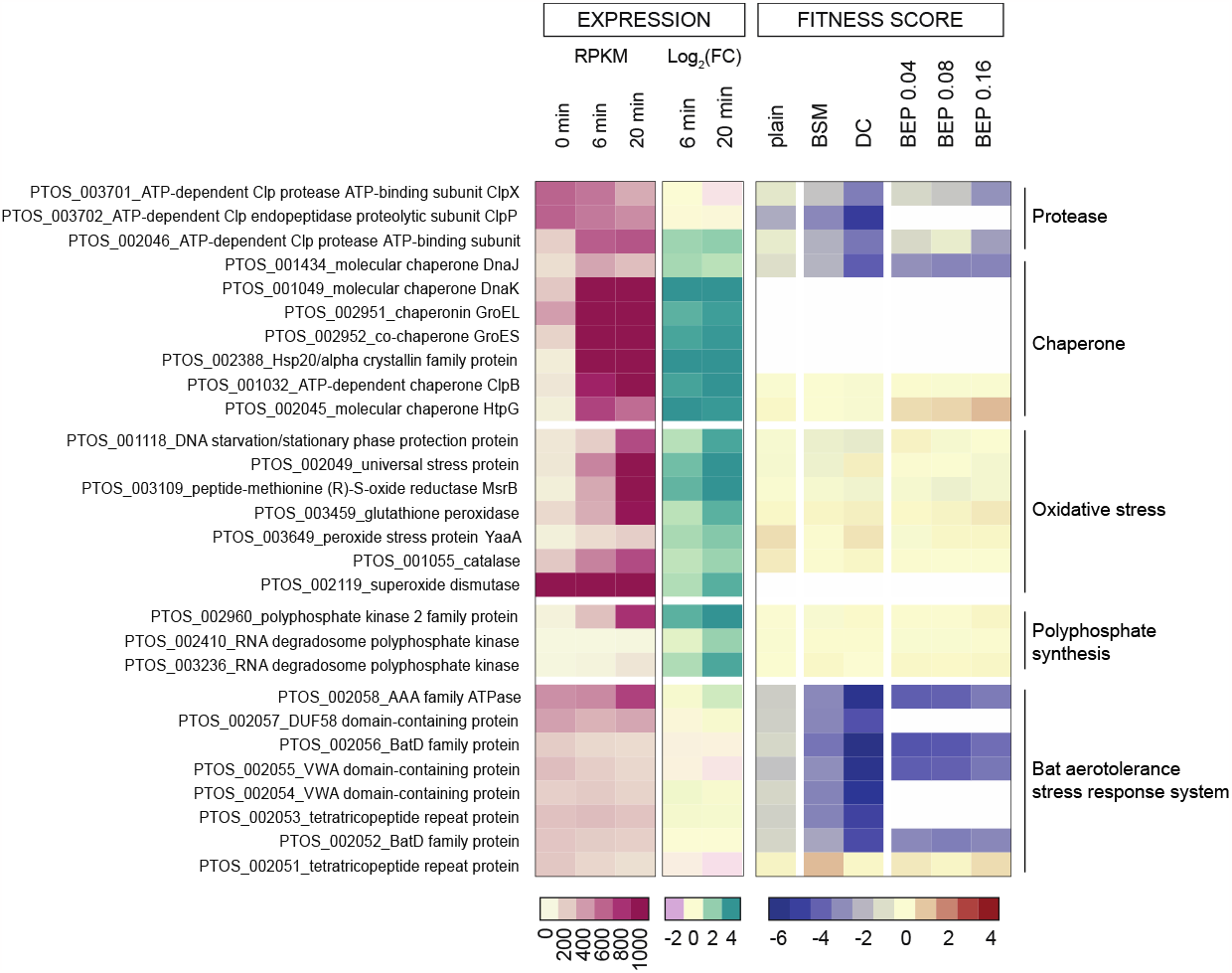
A broad repertoire of stress response systems enable survival in deoxycholate. Heat maps of expression and fitness data for various stress response genes presented as in Figure S7. Protein chaperones, which are important for responding to unfolded proteins or other proteotoxic stress are broadly upregulated in by deoxycholate exposure. The ClpXP protease, which serves a similar function, is not transcriptionally regulated, but is important for fitness. Oxidative stress response genes are broadly upregulated, but no single gene is critical for fitness. Polyphosphate synthesis genes exhibit a similar pattern. The bat system, which has been characterized in aerotolerance, is constitutively expressed and critical for fitness during bile exposure.

