## Supplementary material for "Bile acid fitness determinants of a *Bacteroides fragilis* isolate from a human pouchitis patient": Table S3

**Table S2: Mapping statistics**

| **Category** | **Value** |
| --- | --- |
| Mapped reads | 10,834,060 |
| Barcodes seen 8 or more times | 82,357 |
| Barcodes mapped reliably (minFrac 0.7 minRatio 7)^a^ | 51,383 |
| Barcodes remaining after masking off-by-1 barcodes | 49,543 |
| Distinct locations hit | 43,295 |
| Fraction of reads for those barcodes | 76.6% |
| Insertions in central 10-90% of genes | 37,119 (32310 distinct locations) |
| Protein coding genes with central insertions | 3,489 of 3,967 (87.9%) |
| Strains per hit protein | median 8, mean 10.5 |
| Gene and transposon on same strand | 50.1% |
| Reads per hit protein | median 846, mean 1,456 |
| Reads per million for hit proteins | median 78.1, mean 134.4 |

^a^ minFrac is the minimum fraction of reads for the barcode that agree with the preferred mapping. minRatio is the minimum ratio of reads for preferred mapping over the 2nd-most-frequent mapping.
