## Supplementary material for "Bile acid fitness determinants of a *Bacteroides fragilis* isolate from a human pouchitis patient": Table S7

**Table S7: Strains and primers**

| **Strain** | **Genotype (description)** | **Source** |
| --- | --- | --- |
| AMD776 | E. coli WM3064 / pTGG46_NN1  (Donor strain carrying barcoded ErmR TnHimar containing plasmid) | (1) |
| FC3835 | Bacteroides fragilis str. P207  (Isolate from pouchitis patient with inflammation) | (2) |
|  | B. fragilis str P207 :: pTGG46_NN1  (mixed pool of barcoded TnHimar mutants) | This study |
| **Primer / oligo** | **5’-3’ Sequence (description)** | **Notes** |
| Y adapter oligos | | |
| Mod2_TruSeq | /5'P/GATCGGAAGAGCACACGTCTGAACTCCAGTCA  (5’ phosphorylation) | Y-adapter (3) |
| Mod2_TS_Univ | ACGCTCTTCCGATC*T  (3' T is phosphorothioate bonded) | Y-adapter (3) |
| For two-step nested amplification of barcode junctions - TNseq | | |
| TS_bs_T7-35 | ACACTCTTTCCCTACACGACGCTCTTCCGATCTNNNNNNCGACTCACTATAGGGGATA**GATGTCCACGAG**GTCT  (TruSeqRead1-hexamer-TN specific) | Modified from Nspacer_barseq_universal (3) to increase transposon specific sequence |
| TS_R | GTGACTGGAGTTCAGACGTGTGCTCTTCCGATCT  (TruSeqRead2) |  |
| P5-TS F | AATGATACGGCGACCACCGAGATCTACACTCTTTCCCTACACGACGCTCTTCCGATCT  (P5 sequence-TruSeqRead1) |  |
| P7_MOD_TS_index6 | CAAGCAGAAGACGGCATACGAGAT**CGTGAT**GTGACTGGAGTTCAGACGTGTGCTCTTCCGATCT  (P7 sequence-**index**-TruSeqRead2) | (3) |
| For amplification of barcodes - BarSeq | | |
| Barseq_P1 | AATGATACGGCGACCACCGAGATCTACACTCTTTCCCTACACGACGCTCTTCCGATCTNNNNNGTCGACCTGCAGCGTACG  (P5-TruSeqRead1-hexamer-TN specific) | (3) |
| Barseq_P2_ITxxx | CAAGCAGAAGACGGCATACGAGATXXXXXXGTGACTGGAGTTCAGACGTGTGCTCTTCCGATCTGATGTCCACGAGGTCTCT  (P7 sequence-index-TruSeqRead2-TN specific) | Uniquely indexed primers were used for each sample and are fully listed in (3) |
| For arbitrary nested amplification of barcode junctions in single clones | | |
| U1 F | GATGTCCACGAGGTCTCT |  |
| M13F-N_7_ | TGTAAAACGACGGCCAGTNNNNNNN  (M13F-random heptamer) |  |
| U2 out | CGTACGCTGCAGGTCGAC |  |
| M13F | TGTAAAACGACGGCCAGT |  |
| **For RT-qPCR** | | |
| PTOS_001192_F | GCATCGGGTTGCTAATCGTT | Hypothetical |
| PTOS_001192_R | AGAAGCATCACGAGCCAACA |  |
| PTOS_001365_F | ACACCTCTCCCACCCCAATA | DEAD/DEAH box helicase |
| PTOS_001365_R | GCAGCTGTTTTACCGGTACCT |  |
| PTOS_001272_F | CGAAATTGCAAAGAACACCGGT | Integration host factor |
| PTOS_001272_R | CGAAGCTACCAAATCCACGG |  |
| PTOS_003612_F | CTTGCATTCCTCACCCTGCT | RND efflux transporter |
| PTOS_003612_R | TTCGGAGCTGCGATGACTAC |  |
| PTOS_001049_F | TTGCCCTTGCCCTCTTTTCT | DnaK |
| PTOS_001049_R | CACAGCCACCATAGTAGCGT |  |
| PTOS_002119_F | GGCATTGAGTTGATCGCATCA | Superoxide dismutase |
| PTOS_002119_R | AGTACTGCACTTTTCAGCCA |  |
| PTOS_001852_F | GGCATTGAGTTGATCGCATCA | Serine hydroxymethyltransferase |
| PTOS_001852_R | CACAGCCACCATAGTAGCGT |  |
| PTOS_001202_F | ACAGTGTTCCGTCCTCCAAC | σ^70^ RNA polymerase sigma factor |
| PTOS_001202_R | GCATCGGGTTGCTAATCGTT |  |
